# Developmental programming of hematopoietic stem cell dormancy by evasion of Notch signaling

**DOI:** 10.64898/2026.01.02.697352

**Authors:** Patricia Herrero-Molinero, Eric Cantón, María Maqueda, Cristina Ruiz-Herguido, Arnau Iglesias, Jessica González, Brandon Hadland, Lluis Espinosa, Anna Bigas

**Author notes:** Corresponding author:* Anna Bigas, Cancer Research Program. CIBERONC., Hospital del Mar Research Institute Barcelona (HMRIB), Doctor Aiguader 88, 08003, Spain.

## Abstract

Dormant hematopoietic stem cells (HSCs) are a rare subset of deeply quiescent, metabolically inactive cells that preserve lifelong hematopoietic regeneration. In mice, these cells were thought to arise exclusively after birth in the BM, while fetal liver HSCs were considered uniformly proliferative and rapidly expanding. Here, using H2B-GFP label-retention strategies, we identify a previously unrecognized population of low-dividing HSCs that is specified during embryogenesis and persists into adulthood. We show that this dormant state is established in specific fetal HSCs that evade the activation of Notch signaling within the fetal liver niche, rather than through global loss of pathway activity. Genetic and transient perturbation of Notch signaling reveals a narrow developmental window in during which reduced Notch exposure promotes entry into a durable dormancy program, whereas irreversible pathway disruption compromises long-term function. Notably, transient Notch during fetal development, but not in adult BM, induces HSC quiescence while enhancing long-term repopulation capacity. Together, these findings demonstrate that HSC dormancy is developmentally programmed before birth and identify evasion of Notch signaling as a key mechanism establishing this state, fundamentally revising current models of how long-term regenerative potential is specified.

## INTRODUCTION

Hematopoietic stem cells (HSCs) sustain blood production throughout life by balancing self-renewal and differentiation into all blood lineages. However, HSCs are functionally heterogeneous, with distinct subpopulations arising during development and persisting in adulthood. This includes early lineage-biased subsets, as well as postnatal populations that differ in proliferative status, ranging from actively cycling to deeply quiescent, known as dormant HSCs(Benz, Copley et al. 2012). While epigenetic memory is thought to maintain this diversity(Yu, Yusuf et al. 2017), the mechanisms specifying these states remain unclear.

HSC development occurs in distinct anatomical niches: specification in the aorta-gonad-mesonephros (AGM) region, expansion mainly in the fetal liver (FL), and maintenance in the adult bone marrow (BM)(Dzierzak and Bigas 2018). Although the AGM and BM environments have been extensively characterized, the FL niche remains less understood. FL HSCs undergo robust proliferation, expanding up to 20-fold(Ema and Nakauchi 2000), but recent studies, have identified a slow-dividing subpopulation in the FL that contribute minimally to the hematopoietic system during the first months of life(Ganuza, Hall et al. 2022, Yokomizo, Ideue et al. 2022, Ishida, Mercoli et al. 2025), suggesting that dormancy may arise earlier than previously thought.

Dormant HSCs are characterized by deep quiescence (G0), superior long-term self-renewal, and low metabolic activity. In adult mice, they comprise around 15-30% of the long-term HSC pool and act as a reserve population, activated in response to stress such as infection, blood loss or chemotherapy(Wilson, Laurenti et al. 2008, Trumpp, Essers et al. 2010). While recent work noted a postnatal shift toward quiescence(Bowie, McKnight et al. 2006), rare dormant cells may have been overlooked due to technical limitations. Our prior studies(Thambyrajah, Maqueda et al. 2024), and other recent work(Ganuza, Hall et al. 2022, Ishida, Mercoli et al. 2025) suggest that HSCs with a dormant molecular signature exist during embryogenesis. We previously showed that loss of IκBα perturbs retinoic acid signaling, promoting dormancy in embryonic HSCs (Thambyrajah, Maqueda et al. 2024), highlighting the compatibility of quiescence within the developmental environment, also shown in other systems(Chakkalakal, Christensen et al. 2014, Fuentealba, Rompani et al. 2015)

Notch signaling is essential for the emergence of HSCs from hemogenic endothelium in the AGM region(Kumano, Chiba et al. 2003, Robert-Moreno, Guiu et al. 2008). In contrast, its role in adult hematopoiesis appears limited under steady-state conditions, where Notch signaling is largely dispensable for HSC maintenance(Mancini, Mantei et al. 2005, Maillard, Koch et al. 2008). However, in contexts of regenerative stress, such as post-injury or transplantation, Notch activity has been shown to support HSC expansion and self-renewal(Butler, Nolan et al. 2010, Varnum-Finney, Halasz et al. 2011). By comparison, the role of Notch signaling within the FL niche remains incompletely understood, as genetic disruption of Notch impairs FL HSC function(Gerhardt, Pajcini et al. 2014, Lakhan and Rathinam 2020, Shao, Paik et al. 2023), whereas highly repopulating FL HSCs exhibit low or absent Hes1 expression(Souilhol, Lendinez et al. 2016). In human adult HSCs, downstream of Notch, the transcriptional repressor HES1 reinforces quiescence by maintaining cell cycle arrest(Yu, Alder et al. 2006). Similar Notch-dependent mechanisms govern dormancy in other somatic stem cell systems or in cancer populations(Indraccolo, Minuzzo et al. 2009, Lahmann, Bröhl et al. 2019, Nguyen, Bauer et al. 2019). In the adult skeletal muscle, Notch maintains satellite cells in a quiescent state(Lahmann, Bröhl et al. 2019), and in the adult brain, dynamic oscillations in Hes1 expression regulate the balance between neural stem cell quiescence and activation^(^Sueda, Imayoshi et al. 2019).

In this study, by conditionally deleting RBPJ, the central transcriptional mediator of Notch signaling, or by enforcing overexpression of the Notch intracellular domain (NICD) in the hematopoietic system shortly after HSC emergence using the Vav1-Cre system, we uncover a developmental role for Notch signaling in promoting HSC activation, with reduced Notch exposure permitting entry into dormancy-associated programs. Importantly, our results demonstrate that a specific population of low dividing, label retaining HSCs specified during embryonic development will contribute to the established dormant reservoir of HSCs. Together, our findings demonstrate that dormancy state in HSCs is established in the embryo and Notch signaling plays a previously unrecognized role in balancing HSC dormancy and activation specifically during development, providing new insights into how long-term hematopoietic potential is programmed early in life.

## RESULTS

### Notch-deficient HSCs display increased quiescent and a dormant-like signature but impaired self-renewal

Impairment of Notch signaling in adult BM HSCs has been reported not to affect HSC function across multiple experimental models(Mancini, Mantei et al. 2005, Maillard, Koch et al. 2008, Benveniste, Serra et al. 2014). Nevertheless, by using the Vav1-Cre system, which targets hematopoietic cells after their specification in the embryo(Chen, Yokomizo et al. 2009), to conditionally delete the Notch effector RBPj in the hematopoietic compartment, we observed a significant increase in the subpopulation of quiescent HSCs residing in the G0 phase as previously reported(Lakhan and Rathinam 2020), while the total number of BM Lineage^−^ sca1^+^ckit^+^ (LSK) and HSCs remained comparable to controls (Figure S1A-C). RBPj^vav1^-deficient HSCs exhibit reduced engraftment capacity in primary transplantation assays (Figure S1D) due to a nearly 3-fold reduction in competitive repopulating unit (CRU) frequency compared to controls (Figure S1F) and display a lack of self-renewal capacity in secondary transplantation (Figure S1E). As expected, T cell differentiation was impaired in these mice (Figure S1G-H) whereas multipotent progenitor (MPP) populations remained unaffected (Figure S1I). Notably, transcriptomic profiling of RBPj^vav1^ wild-type (WT) and knockout (KO) BM HSCs (LSK CD150+CD48-) revealed an enrichment of a dormancy-associated gene signature in the RBPj-deficient population (Figure S1J-K).

Given that Vav1-Cre-mediated deletion occurs during embryonic development, we next examined hematopoietic stem and progenitor cells (HSPCs) in the FL at embryonic day 14.5 (E14.5). Flow cytometry analysis showed no significant differences in total LSK or HSPC numbers between RBPj^vav1^ WT and KO embryos (Figure 1A), but a greater proportion of KO HSCs were in the G0 phase (28±7% compared to 48±12%) (Figure 1B). Functional assessment via competitive transplantation of limiting cell doses demonstrated a dramatic reduction in the engraftment potential of RBPj^vav1^-deficient FL HSCs (Figure 1C), consistent with previous findings(Gerhardt, Pajcini et al. 2014, Lakhan and Rathinam 2020). Specifically, we observed a 72-fold reduction in the frequency of repopulating HSCs in RBPj^vav1^ KO compared to WT embryos (Figure 1D).

**Figure 1.**
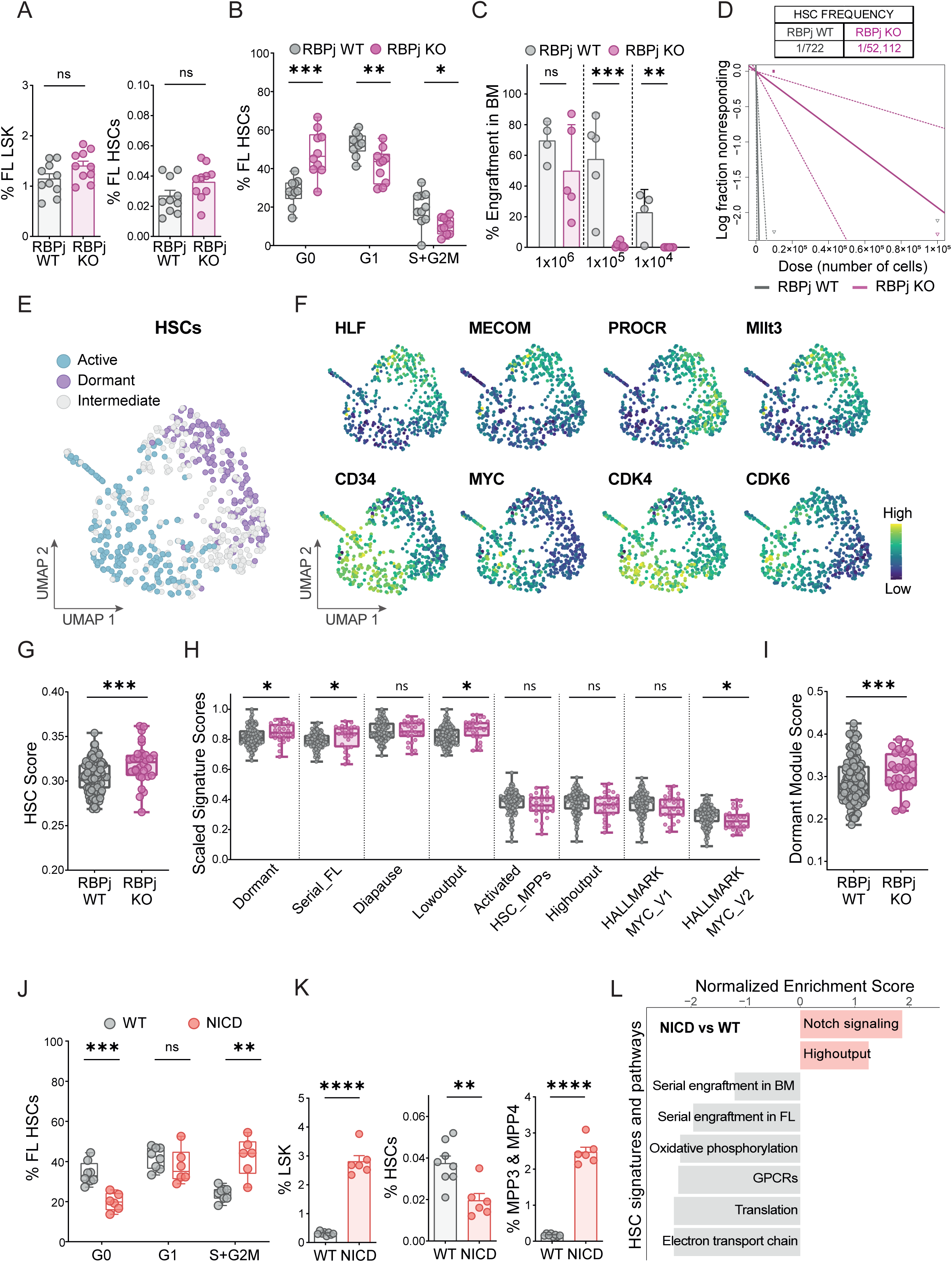
Fetal RBPj^VAV1^ deficiency induces quiescence and a dormancy-associated transcriptome in both FL and BM HSCs but impairs their functionality, whereas NICD^VAV1^ expression promotes FL HSC activation and differentiation. **(A)** Bar plots showing the frequency of LSK (left) and HSCs (LSK CD48^−^ CD150^+^) (right) in RBPj^vav1^ WT and KO E14.5 FL embryos. Each dot represents one embryo (n=10). **(B)** Box plots showing cell cycle analysis of RBPj^vav1^ WT and KO HSCs (LSK CD48^−^ CD150^+^) from E14.5 FL, assessed by Ki67 and DAPI staining. Each dot represents one embryo (n=10). **(C)** Competitive transplantation of the indicated numbers of E14.5 FL cells from RBPj^vav1^ WT and KO mice (CD45.2^+^) together with 2×10^5^ C57BL/6 BM competitor cells (CD45.1^+^) into lethally irradiated recipient mice (CD45.1^+^). Graph shows the percentage of engraftment in BM 16 weeks after primary transplantation. Each dot represents one mouse (n=4-8). **(D)** Limiting dilution assay (LDA) to determine the colony-repopulating unit (CRU) frequency of E14.5 FL HSCs from RBPj^vav1^ WT and KO embryos, analyzed by ELDA (Extreme Limiting Dilution Analysis) at 16 weeks post-transplantation. The table summarizes the estimated HSC frequencies obtained from the LDA. N=4-8 mice. **(E)** UMAP representation of active, intermediate and dormant HSC populations from RBPj^vav1^ WT and KO E14.5 FL LSK scRNA-seq data, based on the Dormant HSC signature and the Active_HSC_MPPs signature. N=2 pooled embryos. **(F)** Expression of HSC-related genes (upper; Hlf, Mecom, Procr, Mllt3) and active HSC-associated genes (bottom; CD34, Myc, Cdk4, Cdk6) in HSC subcluster represented in UMAP. N=2 pooled embryos. **(G-I)** Box plots showing score representation of HSC Score (G); Signature scores including adult HSC dormancy, serial-engrafting FL HSCs, diapause, low-output, activated HSC/MPPs, high-output, MSigDB, Hallmark Myc target genes version 1 (V1) and version 2 (V2) (H); and Dormancy module score (I) in dormant HSC populations from RBPj^vav1^ WT and KO E14.5 FL LSK scRNA-seq data. N=2 pooled embryos. **(J)** Box plots showing cell cycle analysis of WT and NICD^vav1^ HSCs (LSK CD48^−^ CD150^+^) from E14.5 FL. Each dot represents one embryo (n=6). **(K)** Bar plots showing the frequency of LSK (left), HSCs (LSK CD48^−^ CD150^+^) (middle) and MPP3/MPP4 (LSK CD48^+^ CD150^−^) in WT and NICD^vav1^ E14.5 FL. Each dot represents one embryo (n=6). **(L)** Bar plot showing GSEA results from bulk RNA-seq analysis comparing NICD^vav1^ versus WT HSCs (LSK CD48^−^ CD150^+^), using gene signatures previously associated with dormant HSCs, activated HSCs/MPPs, metabolic activity, cell cycle and Notch signaling, including the Notch_signaling hallmark (MSigDB_HALLMARK) as well as signatures for High-output, serially engrafting FL HSCs and serially engrafting BM HSCs. Only statistically significant pathways are shown (adjusted p-value < 0.1). Pathways are sorted in descending order of normalized enrichment score (NES). The experiment included five samples in total (n=3 NICD^vav1^ HSCs and n=2 WT HSCs). Box plots: center line indicates the median; lower and upper hinges represent the first and third quartiles; whiskers extend to the largest and smallest values within 1.5× the interquartile range (IQR). No outliers are present.

To further dissect the molecular differences between RBPj^vav1^ WT and KO HSCs, we performed single-cell RNA sequencing (scRNA-seq) on sorted LSK cells from E14.5 FLs. Thirteen distinct subpopulations were identified by annotating our cells against a reference dataset of mouse E14.5 FL from Gao et al (Gao, Shi et al. 2022). Cell identity distributions are shown using UMAP and PHATE representations (Figure S2A-B), including an HSPC cluster comprising both HSCs and MPPs (Figure S2C-D). Within the HSC cluster, we distinguished dormant and active HSC populations, as well as an intermediate state, based on the expression levels of established HSC dormancy and activation markers (Figure 1E-F and S2E-G), similar to what is found in a recent report(Ishida, Mercoli et al. 2025). In this cluster, we found a similar distribution of WT and RBPj^vav1^ KO cells, nevertheless within the dormant population, RBPj^vav1^ KO HSCs exhibited a significantly higher dormant profile compared to controls (Figure 1G-I), including enrichment of a gene signature previously associated with serial FL transplantation capacity (Figure 1H) and stemness-associated genes (Hlf, Mecom, Procr, Mllt3, Meg3 and H19), as reflected in the Dormancy gene module (Figure 1I and S2F).

Next, we investigated the effect on FL HSCs of constitutive activation of Notch1. To test this, we expressed the Notch intracellular domain (NICD) using the Vav1-Cre system and analyzed E14.5 FL HSCs. We found that NICD^vav1^ HSCs had a decrease in G0-phase (34±6% compared to 20±4%) while increased in cycling phase (S+G2/M) (24±3% compared to 42±10%) (Figure 1J). Moreover, NICD^vav1^ expanded the overall LSK compartment while decreasing the relative proportion of HSCs and promoting their differentiation into MPP3 and MPP4 populations (Figure 1K and S2H). Transcriptome analysis of WT versus NICD^vav1^ E14.5 FL HSCs confirmed Notch pathway activation and downregulation of engraftment-associated signatures (Figure 1L), together with reduced expression of HSC marker genes and upregulation of genes associated with HSC activation (Figure S2I)

Altogether, these results indicate that irreversible disruption of Notch signaling in HSCs during embryonic development induces dormancy-associated transcriptional changes that persist into adulthood, but compromises long-term HSC function, thereby distinguishing this state from physiological dormancy.

### Label-retaining HSCs are generated during embryonic development

Our results suggest the presence of nascent HSCs in the embryo that adopt distinct fates depending on Notch exposure, with cells that evade Notch activation maintaining a low-dividing, dormant state.

To investigate the potential existence of dormant cells during fetal development, embryonic nascent HSCs were labeled with BrdU at E9.5, E10.5 and E11.5, and label retention was analyzed in the FL at E12.5, E14.5 and E16.5, as well as at postnatal days 2 (P2) and 3 (P3) (Figure S3A-F). Despite incomplete labeling of embryonic HSCs (Figure S3B), 2±1% retained high levels of BrdU by P3 in neonatal liver, although lower levels of BrdU labeling in 48±14% P3 HSCs was detected when compared to unstained control (data not shown). Because BrdU incorporation depends on cell division during the labeling window, quiescent or slowly cycling HSCs remained unlabeled, preventing a comprehensive analysis of the entire HSC compartment (only 51±6% E12.5 HSCs were labelled, Figure S3B). Moreover, BrdU-retaining HSCs cannot be functionally characterized, as BrdU detection requires cell fixation and permeabilization.

To overcome these limitations, pTRE-H2B-GFP reporter mice were crossed with ROSA26^M2rtTA^ mice. In this inducible system, doxycycline administration drives H2B-GFP expression, thereby labeling cells. Upon cell division, H2B-GFP is gradually diluted and ultimately lost, enabling the identification of slow-cycling or label-retaining cells (Figure 2A).

**Figure 2.**
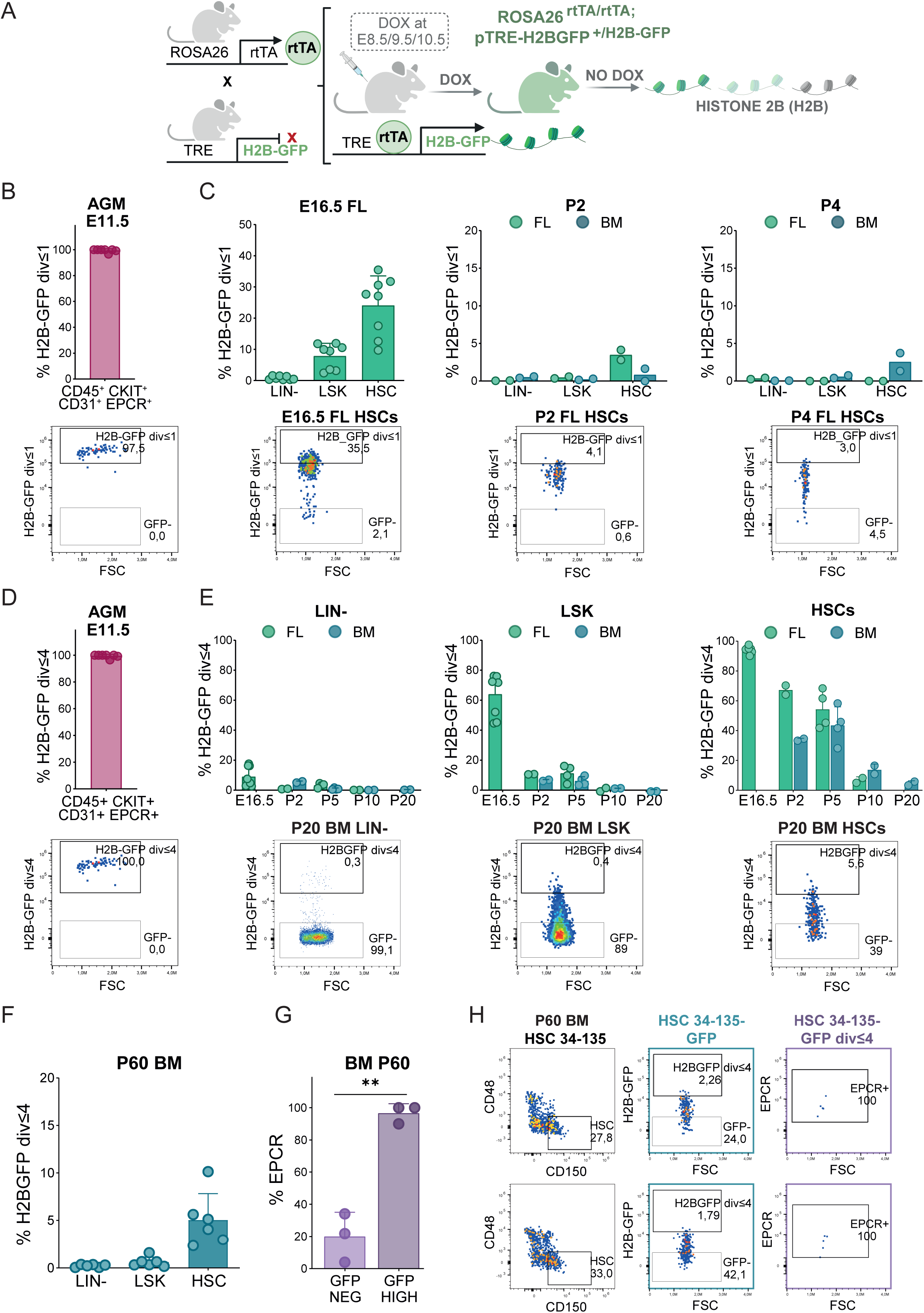
HSCs retaining high GFP levels undergo limited divisions during fetal development. Long-term label-retention assays were performed using ROSA26^M2rtTA/M2rtTA^ mice crossed with pTRE-H2BGFP^+/GFP^ mice. Doxycycline (DOX) administration to pregnant females at E8.5, E9.5, and E10.5 to label cells. **(A)** Experimental scheme. **(B)** Bar plot showing the frequency of cells that do not divide or divide only once (H2B-GFP^div≤1^) within the E11.5 AGM CD45^+^ c-Kit^+^ CD31^+^ EPCR^+^ population (top) and representative dot plot (bottom). Each dot represents one embryo (n=7). **(C)** Bar plots showing the frequency of H2B-GFP^div≤1^ cells in FL (green) and BM (blue) at the indicated developmental stages (E16.5, right; P2, middle; and P4, left) within Lin-, LSK and HSC populations. A representative dot plot from E16.5, P2 or P4 HSCs is shown below. FL HSCs were defined as LSK CD48^−^ CD150^+^, whereas BM HSCs were defined as LSK CD48^−^ CD150^+^ CD34^−^ CD135^−^. Each dot represents one embryo/mouse (n=2-8). **(D)** Bar plot showing the frequency of cells that divide four time or fewer (H2B-GFP^div≤4^) within the E11.5 AGM CD45^+^ c-Kit^+^ CD31^+^ EPCR^+^ population (top) and representative dot plot (bottom). Each dot represents one embryo (n=7). **(E)** Bar plots showing the frequency of H2B-GFP^div≤4^ cells in FL (green) and BM (blue) at the indicated developmental stages (E16.5, P2, P5, P10 and P20) within Lin-(left), LSK (middle) and HSC (right) populations. A representative dot plot from P20 BM Lin-, LSK or HSCs is shown below. FL HSCs were defined as LSK CD48^−^ CD150^+^, whereas BM HSCs were defined as LSK CD48^−^ CD150^+^ CD34^−^ CD135^−^. Each dot represents one embryo/mouse (n=2-8). **(F)** Bar plots showing the frequency of H2B-GFP^div≤4^ cells in P60 BM Lin-, LSK or HSC (LSK CD48^−^ CD150^+^ CD34^−^ CD135^−^) populations. Each dot represents one mouse (n=6). **(G)** Bar plots showing the frequency of EPCR expression in P60 BM HSC (LSK CD48^−^ CD150^+^ CD34^−^ CD135^−^) among H2B-GFP^div≤4^ and GFP-negative cells. Each dot represents one mouse (n=3). **(H)** Representative dot plots of P60 BM HSCs (LSK CD34^−^ CD135^−^ CD48^−^ CD150^+^) showing H2B-GFP^div≤4^ expression and the corresponding EPCR levels. Two biological replicates are shown (replicate 1, top; replicate 2, bottom).

To label nascent HSCs, pregnant females were administrated doxycycline (DOX) on three consecutive embryonic days (E8.5, E9.5, and E10.5). Under these conditions, 99±1% of CD45^+^ CD31^+^ c-Kit^+^ EPCR^+^ cells isolated from dissected E11.5 AGM regions were GFP-labeled (Figure 2B). To estimate the number of HSC cell divisions during fetal and early postnatal development, we applied a GFP dilution-based mathematical model^6^ (Figure S4A). We found that approximately 24±10% of HSCs did not divide or underwent a single division between the E11.5 AGM and the E16.5 FL and these cells were still detected in the neonatal liver at P2 (3±1%) and in the BM at P4 (3±2%) (Figure 2C). Notably, these findings identify a subset of low-dividing HSCs during fetal development.

Subsequently, we monitored H2B-GFP expression over time in distinct hematopoietic populations. HSCs retaining high levels of H2B-GFP, corresponding to four or fewer divisions (Figure S4B), were analyzed at E16.5, P2, P5, P10 and P20 in the FL and/or BM (Figure 2D-E and S4E). We found that the majority of Lin-cells had lost H2B-GFP expression by day E16.5, with 9±7% of cells retaining high GFP intensity. These high-retaining GFP cells were enriched within the LSK population, comprising up to 64±14% at E16.5. This proportion of LSK declined markedly to 11±0.1% by P2, with an almost complete loss observed by P10 (1.2±0.1%). In contrast, a slower gradual decline in H2B-GFP intensity was observed within the HSC population from 94±2% at E16.5 in the FL to 49±13% at P5 in BM and FL and 5±2% at P20 in the BM (Figure 2E). Importantly, a small subset of HSCs with high H2B-GFP retention persisted in the BM that was detectable up to two months of age (P60) (Figure 2F, H) and underwent up to 3.8 divisions (Figure S4C-D). GFP high-retaining P60 BM HSCs expressed superior EPCR levels (Figure 2G-H), which is associated to higher engraftment capacity(Subramaniam, Talkhoncheh et al. 2019, Anjos-Afonso, Buettner et al. 2022).

### The dormancy program in HSCs is established during embryonic development

To assess the engraftment and self-renewal capacity of H2B-GFP high retaining cells, we purified 29 HSCs (LSK CD150^+^ CD48^−^) from P5 neonatal liver and BM, sorted from both the high and low H2B-GFP populations, and transplanted into lethally irradiated recipient mice with 50,000 BM competitor cells (Figure 3A). We observed that high H2B-GFP retaining HSCs from both liver and BM (Figure 3B) exhibited superior repopulation capacity in primary and secondary transplants with higher contribution of LSK and HSCs (Figure 3C-F and S5A-B), consistent with the dormant state ascribed to label-retaining HSCs. In contrast, H2B-GFP low cells failed to exhibit engraftment capacity under these experimental conditions (Figure 3C-F).

**Figure 3.**
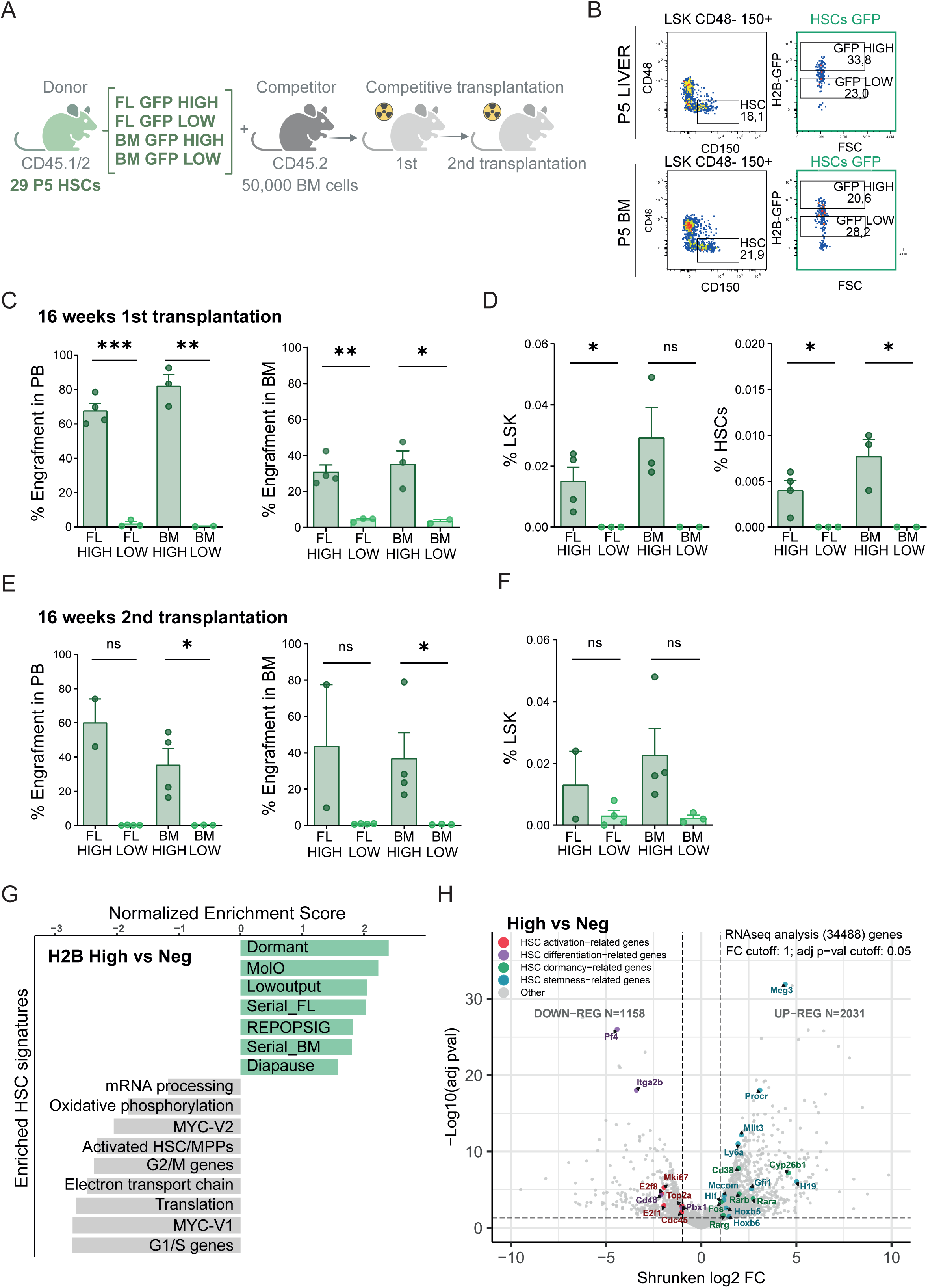
Dormancy is established in a subset of fetal HSCs during embryogenesis. Long-term label-retention assays were performed in pTRE-H2BGFP^+/GFP^;ROSA26^M2rtTA/M2rtTA^ mice, with doxycycline (DOX) administered to pregnant females at E8.5, E9.5, and E10.5. **(A-F)** Following DOX administration at E8.5, E9.5 and 10.5, a total of 29 P5 neonatal liver and BM HSCs (LSK CD48^−^ CD150^+^) were sorted based on H2B-GFP intensity (high and low) and used in a competitive transplantation assay. Specifically, 29 HSCs from P5 liver or BM (CD45.1^+^/CD45.2^+^) with high and low H2B-GFP levels were co-transplanted with 5×10^4^ C57BL/6 (CD45.2^+^) BM competitor cells into lethally irradiated C57BL/6 recipient mice (CD45.2^+^). After 16 weeks following primary transplantation, a total of 1×10^6^ BM cells were transplanted into secondary recipients. **(A)** Experimental scheme. **(B)** Dot plots showing HSCs with high and low H2B-GFP levels in neonatal liver (top) and BM (bottom). **(C)** Bars plots showing the proportion of engraftment in peripheral blood (PB) or BM 16 weeks after primary transplantation. **(D)** Bar plots showing the proportion of LSK and HSCs (LSK CD48^−^ CD150^+^) contribution in primary transplantation after 16 weeks. **(E)** Proportion of engraftment in PB and BM 16 weeks after secondary transplantation. **(F)** Contribution of LSK and HSCs to secondary transplantation at 16 weeks. Each dot represents one mouse (n=2-4). **(G)** GSEA of bulk RNA-seq data comparing P60 BM H2B-GFP high versus negative HSCs (LSK CD48^−^ CD150^+^ CD34^−^ CD135^−^), using gene signatures associated with dormant HSCs and activated HSCs/MPPs, metabolic activity and cell cycle, including Dormant HSC, MolO, Low-output, Diapause, serially engrafting FL and BM HSCs, Repopulating, and Active HSCs/MPPs signatures. Only statistically significant pathways are shown (adjusted p-value < 0.1). Pathways are sorted in descending order of normalized enrichment score (NES). The experiment included six samples in total (n=3 GFP high HSCs and n=2 GFP negative HSCs). **(H)** Volcano plot of bulk RNA-seq analysis comparing P60 BM H2B-GFP high and negative HSCs, highlighting stemness-related genes (blue), HSC dormancy-associated genes (green), HSC activation markers (red) and HSC differentiation-related genes (purple). Differentially expressed genes were defined by an FDR adjusted p-value < 0.05 and a shrunken Fold-change (FC) > 1.

To further investigate the nature of H2B-GFP high-retaining HSCs, LSK CD48^−^ CD150^+^ CD34^−^ CD135^−^, exhibiting high or negative levels of H2B-GFP, were sorted for transcriptomic analysis (Figure S5C). We observed that HSCs with high H2B-GFP levels exhibit enrichment of dormancy-associated transcriptional programs, including signatures linked to superior engraftment capacity and stemness, alongside downregulation of gene sets associated with HSC activation (Figure 3G-H and S5D-E). Specifically, H2B-GFP high HSCs express higher levels of retinoic acid related genes, a known regulator of dormancy in adult HSCs (Cabezas-Wallscheid, Buettner et al. 2017) (Figure 3H and S5E). Together, these findings demonstrate that a population of dormant HSCs is established during embryonic development.

### Transient inhibition of Notch in FL HSCs induces a functional dormant phenotype, but not in adult BM HSCs

Irreversible Notch signaling alteration in FL HSCs affect the regulation of the dormancy genetic program, but disrupt their functionality. Based on this, we hypothesized that transient inhibition of Notch signaling in HSCs could induce a dormant functional phenotype. To test this hypothesis, Lin⁻ cells from E14.5 FL were purified and cultured for 48 hours in the presence of either DMSO (control) or γ-secretase inhibitors, DAPT and Compound E (CPDE) which prevent Notch cleavage (Figure 4A). After 48h, we quantified the proportion of FL HSCs and analyzed their cell cycle profiles. Our results showed a significant increase in the percentage of FL HSCs upon Notch inhibition with 1.7- and 2.3-fold increase with DAPT and CPDE, respectively (Figure 4C, S5F and S6B). Additionally, there was a marked increase in the fraction of HSCs residing in a quiescent state, with the percentage of cells in G0 phase rising from 20±5% to 42±9% (Figure 4A-B and S5F). Similar results were observed using specific blocking antibodies against Notch1 and Notch2 receptors, further confirming that the observed phenotype was due to Notch inhibition (Figure S5G).

**Figure 4.**
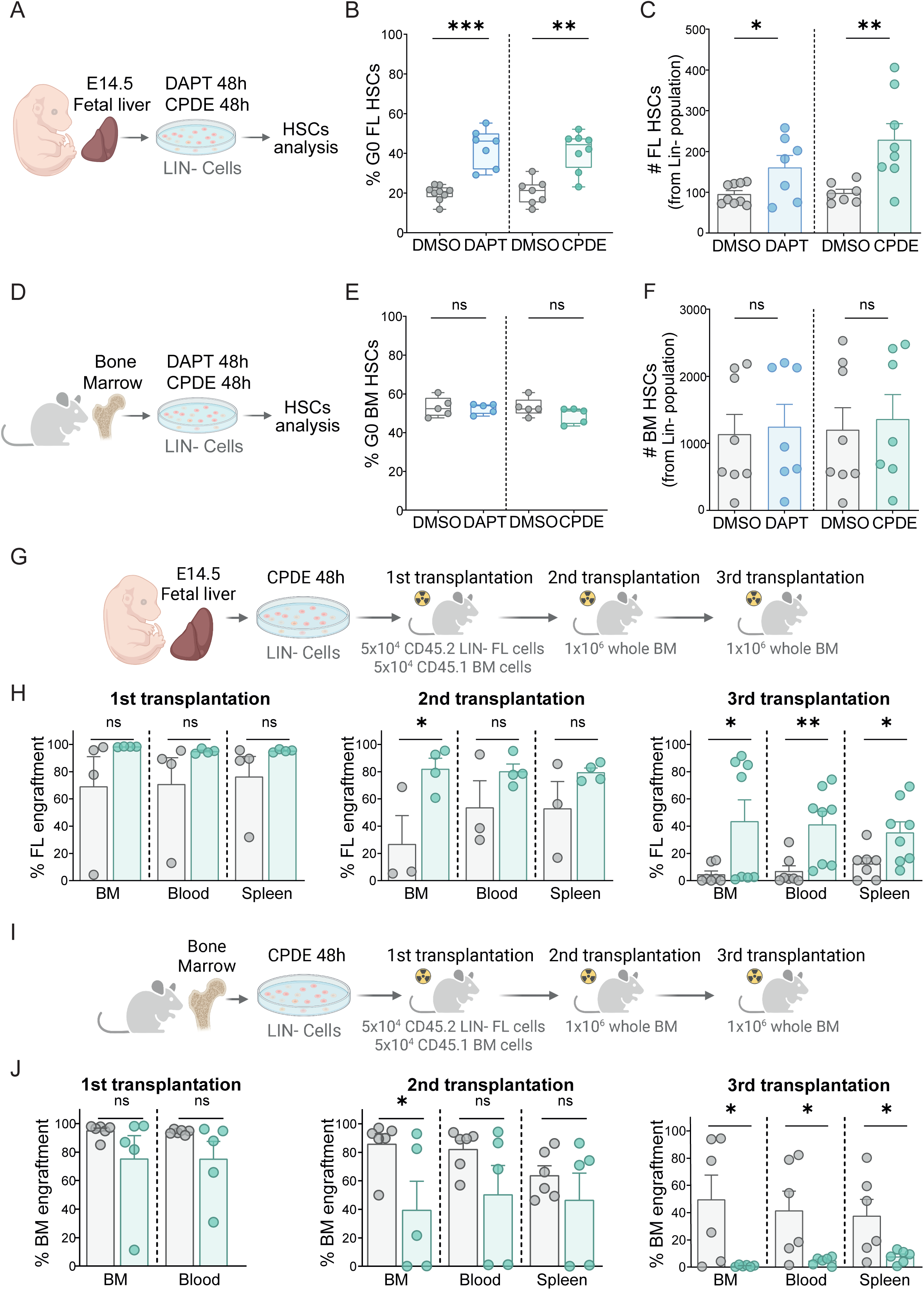
Transient Notch inhibition induces quiescence and preserves FL HSC frequency in vitro, enhancing their self-renewal capacity in vivo, whereas this effect does not occur in adult BM HSCs. Lineage-negative (Lin-; defined by CD3^−^, B220^−^, Ter119^−^ and Gr1^−^) cells from E14.5 FL (A-C, G-H) or BM (D-F, I-J) were culture in vitro for 48 hours with DMSO (control), gamma-secretase inhibitors DAPT (10μM) and Compound E (CPDE, 1μM) to inhibit Notch signaling. **(A, D)** Experimental scheme for transient in vitro Notch inhibition of E14.5 FL (A) and BM (D) cells. **(B, E)** Box plots showing the frequency of G0 control (DMSO) or Notch-inhibited HSCs (LSK CD48⁻ CD150⁺) from E14.5 FL (B) and BM (E), as determined by cell cycle analysis, following treatment with the γ-secretase inhibitors DAPT and Compound E. Each dot represents one embryo or mouse (n=5-9). **(C, F)** Bar plots showing the number of control (DMSO) or Notch-inhibited HSCs in E14.5 FL (C) and BM (F), normalized per 10^5^ Lin-cells. Each dot represents one embryo or mouse (n=5-9). **(G-J)** A total of 50,000 CD45.2^+^ E14.5 FL Lin-(G-H) or BM (I-J) cells treated with DMSO or CPDE for 48 hours were competitively transplanted together with 50,000 CD45.1^+^ BM competitor cells into lethally irradiated CD45.1^+^ C57BL/6 recipient mice. After 16 weeks following primary transplantation, a total of 1×10^6^ BM cells were transplanted into secondary and tertiary recipients. **(G, I)** Experimental scheme of E14.5 FL (G) and BM (I) cells. **(H, J)** Bar plots showing engraftment frequencies in primary (left), secondary (middle) and tertiary (right) transplantation assay (16 weeks each) in peripheral blood, BM and spleen of FL (H) and BM (J) cells.

Next, we investigated the effects of Notch inhibition in adult BM HSCs by culturing Lin-cells in the presence of γ-secretase inhibitors (Figure 4D). Importantly, Notch inhibition did not affect the cell cycle profile of BM HSCs compared to control (Figure 4E-F); however, in BM, longer inhibition of Notch promoted differentiation, leading to decrease in HSCs, LSK and Lin^−^ populations and a concurrent increase in the Lin+ populations (Figure S6A), consistent with previous reports(Butler, Nolan et al. 2010).

Gene expression analysis of γ-secretase inhibitor-treated FL cells confirmed effective Notch pathway suppression, with a reduction of at least 50% in the expression of canonical Notch target genes such as *Hes1, Hey1* and *Hey2* in FL Lin-cells (Figure S6C). Additionally, expression of the activation-associated genes *Cdk6* and *Myc* was markedly reduced in Notch-inhibited E14.5 FL HSCs, supporting their low-dividing state (Figure S6D) and consistent with a dormant phenotype. We next investigated whether Notch-inhibited FL HSCs exhibited functional dormancy. To investigate whether Notch-inhibited FL HSCs exhibited a functional dormant phenotype, we performed serial transplantation experiments using E14.5 FL Lin⁻ cells treated with DMSO (control) or CPDE for 48 hours. In primary transplants, 50,000 CD45.2 treated FL Lin-cells were transplanted with 50,000 CD45.1 BM competitor cells into lethally irradiated CD45.1 recipient mice. Secondary and tertiary transplants were performed four months later with 10^6^ BM cells (Figure 4G). Mice receiving CPDE-treated FL cells showed consistently higher levels of hematopoietic reconstitution with multilineage engraftment in secondary and tertiary transplants, indicating a dormant HSC phenotype (Figure 4H and S6F). Additionally, serial transplantation revealed greater contributions of FL LSK and HSCs (Figure S6E), along with multilineage engraftment similar to DMSO-treated control cells (Figure S6F). In line with our previous findings, we observed a slower rate of single-cell division in vitro (Figure S6G).

In contrast, as suggested by previous genetic deletion studies(Mancini, Mantei et al. 2005, Maillard, Koch et al. 2008, Benveniste, Serra et al. 2014), Notch inhibition in adult BM HSCs did not replicate the effects observed in E14.5 FL HSCs in equivalent serial transplantation experiments (Figure 4I). CPDE-treated BM cells exhibited lower engraftment levels after secondary and tertiary transplants (Figure 4J) and contributed less to the LSK and HSC populations (Figure S6H), with biased myeloid multilineage engraftment (Figure S6I).

Together, these findings, along with genetic models of Notch mutants indicate that Notch signaling imposes an active HSC phenotype. In its absence, HSCs acquired a dormant state as early as FL development.

## DISCUSSION

Through the use of various Notch-mutant mouse models, *in vitro* modulation of Notch and label-retaining strategies, we have demonstrated the existence of low-dividing HSCs in the mouse embryo that require Notch inactivation and that contribute to the dormant adult population of HSCs.

Dormancy is an evolutionarily conserved mechanism that preserves somatic stem cell integrity by shielding them from genotoxic stressors such as mutations, infections, and chemotherapy, thereby maintaining HSC long-term self-renewal and preventing premature differentiation or exhaustion (Passegué, Wagers et al. 2005, Trumpp, Essers et al. 2010). During development, HSCs are exposed to drastic environmental changes from their formation in the AGM, migration to the FL through circulation in extraembryonic tissues and subsequent colonization of the BM. Mechanisms such as dormancy to protect the integrity of HSCs in these changing environments may provide an evolutionary advantage to preserve lifelong hematopoietic potential. We provide evidence that a subset of HSCs can enter a dormant state during embryonic development, contributing to the dormant HSC pool observed in the adulthood.

Label-retaining cell (LRC) strategies have been widely used to detect dormant somatic stem cells(Tumbar, Guasch et al. 2004) including HSCs(Wilson, Laurenti et al. 2008). By using the SCL-tTA-S2 line crossed with H2B-GFP, a dormant population of rare dividing cells (once every 145 days) adult BM HSCs were identified using a Tet-OFF strategy(Wilson, Laurenti et al. 2008). However, the percentage of dormant HSCs is hard to define since it depends on the threshold of GFP and the number of estimated divisions. Our experiments have used a tet-ON model (M2rtTA;H2B-GFP) in which 99.4±1% of CD45^+^EPCR^+^CD31^+^ckit^+^ E11.5 AGM cells are labelled in response to doxycycline. We established the threshold conditions by calculating the number of cell divisions in vivo and the leakage in the H2B-GFP line. We have established two different thresholds for low-dividing HSCs, one gate comprising cells that have divided 0 or once, and high GFP cells comprising cells that have divided less than 4 times. Due to the different strategies used with the BM experiments, the variability of the H2B-GFP system and the embryonic nature of our analysis, it is difficult to establish the total amount of cells labelled in the embryo that may contribute to the previously described adult dormant HSC population. However, we demonstrate that the few high GFP HSCs remaining at 2 months express a dormant signature and are functionally highly repopulating HSCs, making them bona-fide dormant HSCs that were specified during embryonic development. Indeed, the embryonic establishment of dormant somatic stem cells has already been shown for neural stem cells(Fuentealba, Rompani et al. 2015) and likely for muscle stem cells(Chakkalakal, Christensen et al. 2014).

Notch is a known regulator of dormancy in different somatic stem cells(Lahmann, Bröhl et al. 2019), (Sueda, Imayoshi et al. 2019), although the neural and muscle stem cells require Notch activity and Hes1 to maintain the dormant state. However, nothing is known about the role of Notch in establishing the dormant program during embryonic development. After demonstrating that dormant HSCs are specified in the embryo, we identify the Notch pathway as a critical regulator of this process. However, in this context, dormant HSCs arise in a developmental context characterized by reduced or absent Notch activation during their residence in the FL rather than by global pathway inactivation. Our results show that hematopoietic deletion of the Notch effector RBPj after HSC generation induces a quiescent state and a dormancy-associated transcriptome in both FL and BM HSCs, however these cells have impaired engraftment, consistent with previous reports of Notch deficiency in the FL(Gerhardt, Pajcini et al. 2014, Lakhan and Rathinam 2020), but opposite to the lack of effect of Notch deficiency in the adult BM. Our results indicate that the effects of Notch signaling deficiency on fetal HSCs persist into adulthood, ultimately compromising the long-term function of BM HSCs. Using the Vav1-Cre system bypasses the requirement of Notch for HSC generation in the AGM region(Chen, Yokomizo et al. 2009, Ruiz-Herguido, Guiu et al. 2012) but impairs HSC reconstitution capacity under stress conditions such as transplantation, 5-FU treatment and cytokine-induced stress(Gerhardt, Pajcini et al. 2014, Lakhan and Rathinam 2020). Collectively, these findings suggest that the influence of Notch signalling on HSC function is temporally restricted, impacting HSC activity only when modulation occurs during embryonic development, a critical window for fetal HSC maturation and proliferation. In contrast, constitutive activation of Notch signaling through NICD^VAV^ overexpression in FL HSCs promotes their proliferation and differentiation commitment toward MPPs, and finally their leukemic transformation(Gekas, D’Altri et al. 2016).

Given that both permanent deletion of RBPj and constitutive activation of Notch signaling impair the functional capacity of FL and BM HSCs, we investigated whether transient Notch inhibition could modulate the cell cycle status of FL HSCs while preserving their functional potential. Notably, transient Notch inhibition induced quiescence and preserved the overall frequency of FL HSCs *in vitro,* while enhancing their long-term self-renewal capacity and maintaining multilineage differentiation potential *in vivo.* In contrast, this effect was not observed in BM HSCs, further confirming the different usage of Notch in the embryo and in the adult HSCs(Mancini, Mantei et al. 2005, Maillard, Koch et al. 2008, Gerhardt, Pajcini et al. 2014).

In light of the growing evidence that HSC expansion in the FL is limited(Ganuza, Hall et al. 2022), and based on our findings demonstrating that Notch signalling regulates the active versus dormant state of FL HSCs, we propose that a subset of HSCs may enter a dormant state already during embryogenesis. This challenges the prevailing view that dormancy is established only after postnatal week 3 in the BM(Bowie, McKnight et al. 2006).

Our findings raise relevant translational implications for the generation of functional iPSC-derived HSCs. Current differentiation protocols predominantly generate hematopoietic progenitors with limited self-renewal and engraftment capacity upon secondary transplantation, along with transcriptomic profiles resembling HSPCs from the CS14 embryonic AGM region(Ng, Azzola et al. 2016). We speculate that iPSC-derived HSCs may require additional maturation, reflecting developmental processes in the FL niche, to achieve robust long-term functionality. Our findings suggest that transient modulation of Notch signalling during in vitro differentiation may be worth exploring as a strategy to enhance long-term HSC potential.

## DATA AVAILABILITY

Bulk RNA-seq and single-cell RNA-seq datasets generated in this study have been deposited in ArrayExpress/SRA repository under the following accession numbers: E-MTAB-15872 (Bulk RNA-seq BM), E-MTAB-15873 (Bulk RNAseq FL CompE), E-MTAB-15912 (Bulk RNAseq FL NICD) and E-MTAB-15875 (scRNA-seq FL RBPJ^vav1^ KO).

## CODE AVAILABILITY

This study did not involve the development of software. Nevertheless, scripts used to process the generated bulk and single cell RNA-seq datasets are available in Github repository: https://github.com/BigaSpinosaLab/PAPER_Notch_dormancy_fetal_hematopoiesis

## ACKNOWLEDGMENTS

The authors acknowledge all members of Espinosa and Bigas laboratories for helpful discussions and technical support. We also thank the Centre for Genomic Regulation, Universitat Pompeu Fabra Flow Cytometry Unit, animal facility of the PRBB and the Single Cell Unit of the Josep Carreras Leukemia Research Institute for their technical support. All illustrations in the manuscript were created with BioRender.com. This work was supported by Agencia Estatal de Investigación(AEI)/MCIN/10.13039/501100011033, grant PID2019-104695RB-I00, José Carreras Leukäemie-Stiftung, AGAUR, Generalitat de Catalunya (2021SGR39) and ERC synergy 101167065 ‘MakingBlood’ to AB. P.H.M. is a recipient of a Formación de Personal Investigador PhD fellowship (PRE2020-095037) from the Spanish Ministry of Science, Innovation and Universities. M.M. is a recipient of a grant from the Instituto Carlos III (grant number CA22/00011; cofunded by the European Social Fund Plus and by the European Union).

## AUTHOR CONTRIBUTIONS

Contribution: A.B., L.E., and P.H.M. designed the study and experimental design; P.H.M., C.R.H., A.I., and J.G. performed experiments; M.M. and E.C performed bioinformatics analysis; P.H.M., B.H. contributed with data analysis and A.B. and P.H.M. wrote the manuscript with input from all authors; A.B. provided resources and supervised the research.

## SUPLEMENTARY FIGURE TITLES AND LEGENDS

**Figure S1. RBPj^vav1^-deficient BM HSCs are more quiescent than Notch-competent HSCs, display a dormant HSC signature but exhibit a dysfunctional engraftment capacity. Related to Figure 1**.

**(A-B)** Box plots showing cycle analysis of RBPj^vav1^ WT and KO HSCs (LSK CD48^−^ CD150^+^) from adult BM using Ki67 and DAPI staining (A), with representative dot plots (B). Each dot represents one mouse (n=6-11).

**(I) (C)** Bar plots showing the numbers of LSK (left) and HSCs (LSK CD48^−^ CD150^+^, right) per 10,000 BM cells in RBPj^vav1^ WT and KO BM mice. Each dot represents one mouse (n=12-19).

**(D-E)** Competitive transplantation of decreasing doses of adult BM cells from RBPj^vav1^ WT and KO mice (CD45.2^+^), transplanted together with 2×10^5^ C57BL/6 BM competitor cells (CD45.1^+^) into lethally irradiated recipient mice (CD45.1^+^). Bar plots show the engraftment frequencies of RBPj^vav1^ WT and KO BM cells in the BM 16 weeks after primary (D) and secondary transplantation

(E). Each dot represents one mouse (n=4-15).

**(F)** Limiting dilution assay (LDA) to determine CRU frequency by ELDA of adult BM HSCs from RBPj^vav1^ WT and KO mice at 16 weeks post-transplantation. Table summarizing estimated HSC frequencies from LDA (top). Each dot represents one mouse (n=4-15).

**(G)** Percentages of T cells (CD3^+^), B cells (B220^+^ or CD45R^+^), myeloid cells (Mac1^+^ or CD11b^+^) and erythroid cells (Ter119^+^) in the BM of RBPj^vav1^ WT and KO mice. Each dot represents one mouse (n=26-32).

**(H)** Percentages of helper T cells (CD4^+^ CD8^−^), cytotoxic T cells (CD4^−^ CD8^+^) and double-positive thymocytes (CD4^+^ CD8^+^) in the thymus of RBPj^vav1^ WT and KO mice. N= 4 to 14 mice.

**(I)** Percentages of adult BM multipotent progenitors (MPPs): MPP1 (Multipotent; LSK CD34^+^ CD135^−^ CD48^−^ CD150^+^), MPP2 (Megakaryocyte/Erythroid biased; LSK CD34^+^ CD135^−^ CD48^+^ CD150^+^), MPP3 (Myeloid biased; LSK CD34^+^ CD135^−^ CD48^+^ CD150^−^) and MPP4 (Lymphoid biased; LSK CD34^+^ CD135^+^ CD48^+^ CD150^−^).

**(J)** Principal component analysis (PCA) based on the top 500 expressed genes from bulk RNA-seq of 300 RBPj^vav1^ WT and KO BM HSCs (LSK CD48^−^ CD150^+^). The experiment included seven samples in total (n=3 RBPj^vav1^ KO HSCs and n=4 WT BM HSCs).

**(K)** Bar plot displaying GSEA results from bulk RNA-seq RBPj^vav1^ KO versus WT BM HSCs (LSK CD48^−^CD150^+^) and against gene signatures previously associated to dormant HSCs, activated HSCs/MPPs, metabolic activity and cell cycle, including adult HSCs dormancy, WP_GPCRs from Wikipathways (WP), High-output and Multilineage signatures. Only statistically significant pathways are shown (adjusted p-value < 0.1). Pathways are sorted in decreased order by corresponding normalized enrichment score (NES). Experiment included seven samples in total (n=3 RBPj^vav1^ KO HSCs and n=4 WT BM HSCs).

**Figure S2. Single-cell RNA-seq analysis of RBPj WT and KO E14.5 FL HSCs. Related to Figure 1**.

scRNA-seq analysis was performed on FACS-sorted LSK cells from E14.5 FL of RBPj^vav1^ WT and KO embryos. N= 2 pooled embryos.

**(A-B)** UMAP (A) and PHATE (B) representations illustrate the distribution of distinct hematopoietic clusters within the LSK compartment. Cluster annotation was based on the cellular identities defined from Gao et al. (2022). CLP: Common lymphoid progenitors. CMP: Common myeloid progenitors. CP: Common progenitors. MegE: Megakaryocyte and erythrocyte. Mk: Megakaryocytes.

**(C)** UMAP visualizations of HSC/MPP cluster cells colored by RBPj^vav1^ WT and KO genotype (top) or HSC and MPP cell identities (bottom).

**(D)** UMAP visualizations of HSC/MPP cluster cells colored by Stem Score (left) and Active HSC/MPPs signature (right).

**(E)** UMAP projections of HSC cluster cells colored by expression levels of fetal HSC-related genes (H19, Meg3) and active HSC-associated genes (Mki67, Cdk1).

**(F)** Heatmap showing genes comprising the dormancy and activity gene modules and their expression across active, dormant, and intermediate HSC subclusters. Samples are annotated by genotype (RBPj^vav1^ WT and KO), hierarchical clustering, and cell-cycle stage.

**(G)** Heatmap of gene-set scores across active, dormant, and intermediate HSC subclusters, including signatures associated with adult HSC dormancy, serially engrafting FL HSCs, diapause, Low-output, activated HSC/MPPs, High-output, Hallmark Myc target genes version 1 (V1) and version 2 (V2). Samples are annotated by genotype (RBPj^vav1^ WT and KO), hierarchical clustering, and cell-cycle stage.

**(H)** Representative dot plots of E14.5 FL LSK cells (left), HSC/MPPs populations (right) and HSCs (LSK CD48^−^ CD150^+^) cell cycle profiles (bottom) in WT and NICD embryos.

**(I)** Volcano plot from bulk RNA-seq data analysis comparing NICD against WT E14.5 FL HSCs (LSK CD48^−^ CD150^+^), highlighting Notch target genes (pink), HSC activation markers (orange), and HSC markers (blue). Differentially expressed genes were defined by FDR adjusted p-value < 0.1 and shrunken Fold-change (FC) > 0.5.

**Figure S3. BRDU-retaining cells reveal the presence of low-dividing HSCs during fetal development. Related to Figure 2**.

Long-term label-retention assays were performed by administering BrdU to pregnant mice at embryonic days E9.5, E10.5, and E11.5, allowing incorporation of BrdU into the DNA. After a chase period without BrdU, BrdU levels progressively decreased with each cell division.

**(A)** Experimental scheme of BrdU administration (left) and schematic representation of the BrdU label-retaining assay (right).

**(B)** Percentage of BrdU⁺ cells in E12.5 FL populations including Lin⁻ (lineage-negative; CD3⁻, B220⁻, Gr1⁻, Ter119⁻), LSK and HSCs (LSK CD48⁻ CD150⁺). Each dot represents pooled embryos (n= 5).

**(C)** Representative dot plot (left) and histogram (right) of E14.5 FL LSK cells showing BrdU intensity levels used to define BrdU negative, positive and high gating thresholds.

**(D)** Bar plots showing the frequency of BrdU high label-retaining cells at the indicated developmental stages (E12.5, E14.5, E16.5, P2 and P3) within Lin-(left), LSK (middle) and HSC (LSK CD48⁻ CD150⁺; right) populations.

**(E)** Dot plots showing the flow cytometry gating strategy used to identify Lin-, LSK, HSCs and EPCR+ HSCs (top), with corresponding BrdU high levels in each population (bottom).

**Figure S4. H2B-GFP;M2rtTA mouse model and estimation of HSC division rates. Related to Figure 2**.

Following doxycycline (DOX) administration at E8.5, E9.5 and 10.5, FL and BM cells from H2B-GFP;M2rtTA mice were analyzed at the indicated developmental stages (E16.5, P2, P5, P10 and P20) to assess H2B-GFP label-retaining cells.

**(A-B)** Estimation of the actual number of divisions (Na) calculated using the GFP-dilution mathematical model described by Ganuza et al., 2022, based on flow cytometry gating of H2B-GFP levels. Representative dot plots show the mean ± SD of H2B-GFP mean fluorescence intensity (MFI) in cells from the E11.5 AGM (left) and E16.5 FL (right) that did not divide or underwent only one division (Gate for Div≤ 1) (A) or fewer than four divisions (Gate for Div ≤ 4)

(B). E11.5 AGM cells were defined as CD45^+^EPCR^+^CD31^+^cKit^+^ cells and FL HSCs were defined as LSK CD48^−^ CD150^+^ cells.

**(C-D)** Estimation of HSC division rates based on H2B-GFP MFI using the GFP-dilution mathematical model described by Ganuza et al., 2022. (C) Table showing the estimated actual number (Na) of divisions in H2B-GFP retaining HSCs (GFP+) and in HSCs that underwent fewer than four divisions (GFP div≤ 4) from E11.5 AGM through different developmental stages (E16.5,

P2, P5, P10, P20, P60) in the FL and BM. (D) Graphical representation of the estimated Na divisions of HSCs that underwent fewer than four divisions (H2B-GFP div≤ 4) from E11.5 AGM to the FL (left) and BM (right) across developmental stages. FL HSCs were defined as LSK CD48^−^ CD150^+^ cells, whereas BM HSCs were defined as LSK CD48^−^ CD150^+^ CD34^−^ CD135^−^ cells.

**(E)** Representative spectral flow cytometry gating strategy for the identification of hematopoietic populations and analysis of their corresponding H2B-GFP levels in BM at P20. Populations include Lin-cells (lineage-negative; CD3^−^, B220^−^, Gr1^−^, Ter119^−^), LSK cells, HSCs (LSK CD34^−^ CD135^−^ CD48^−^ CD150^+^), MPP1 (LSK CD34^+^ CD135^−^ CD48^−^ CD150^+^), EPCR^+^ HSCs, MPP2 (LSK CD34^+^ CD135^−^ CD48^+^ CD150^+^), MPP3 (LSK CD34^+^ CD135^−^ CD48^+^ CD150^−^) and MPP4 (LSK CD34^+^ CD135^+^ CD48^+^ CD150^−^).

**Figure S5. A subpopulation of HSCs retaining high GFP levels acquire dormancy during embryogenesis. Related to Figure 3**.

Long-term label-retention assays were performed in pTRE-H2BGFP^+/GFP^;ROSA26^M2rtTA/M2rtTA^ mice, with doxycycline (DOX) administered to pregnant females at E8.5, E9.5, and E10.5.

**(A-B)** Following DOX administration at E8.5, E9.5, and E10.5, 29 P5 neonatal liver and BM HSCs (LSK CD48⁻ CD150⁺) were sorted based on high or low H2B-GFP expression and co-transplanted with 5 × 10⁴ C57BL/6 (CD45.2⁺) BM competitor cells in a competitive transplantation assay. Bars plots show the contribution to peripheral blood (left) and BM (right) lineages, including myeloid (Ly6C/Ly6G^+^ or Gr1^+^), erythroid (Ter119^+^), B (B220^+^ or CD45R^+^), and T (CD3^+^) cells, 16 weeks after primary (A) and secondary (B) transplantation. Each dot represents one mouse (n=2-4).

**(C)** Representative spectral flow cytometry gating strategy for sorting P60 BM HSCs (LSK CD34^−^ CD135^−^ CD48^−^ CD150^+^) with high H2B-GFP retention, indicating HSCs that have undergone fewer than four divisions (H2B-GFP div≤ 4), or with negative H2B-GFP levels.

**(D)** Principal component analysis (PCA) of bulk RNA-seq data from 300 P60 BM HSCs (LSK CD34^−^ CD135^−^ CD48^−^ CD150^+^) with high or negative H2B-GFP levels. The experiment included three biological replicates per condition.

**(E)** GSEA of bulk RNA-seq data comparing P60 BM H2B-GFP high versus negative HSCs (LSK CD48^−^ CD150^+^ CD34^−^ CD135^−^), using pathways from WikiPathways. Only statistically significant pathways are shown (adjusted p-value < 0.1). Pathways are sorted in descending order of normalized enrichment score (NES). The experiment included six samples in total (n=3 GFP high HSCs and n=2 GFP negative HSCs).

**(F)** Representative FACS plots showing LSK cells (left), HSCs (LSK CD48^−^ CD150^+^; middle) and cell cycle distribution (right) of HSCs from E14.5 FL embryos treated with DMSO (control, top) or Compound E (bottom) for 48 hours to inhibit Notch signaling.

**(G)** Cell cycle analysis of E14.5 FL HSCs (LSK CD48⁻ CD150⁺) treated for 48 hours with specific Notch receptor–blocking antibodies to inhibit Notch signaling or with IgG as controls (C). Each dot represents one embryo (n=6).

**Figure S6. Transient Notch inhibition in vitro in FL and adult BM HSCs. Related to Figure 4**.

**(A-B)** Representative dot plots of Lin-, LSK and HSCs (LSK CD48^−^ CD150^+^) from adult BM (A) and E14.5 FL (B) treated in vitro with DMSO (control) or DAPT (50μM) to inhibit Notch signaling for 7 days and 72 hours, respectively. Experimental schemes are shown at the top. N= 2 biological replicates.

**(C-D)** Gene expression fold-change measured by quantitative PCR (qPCR) of Notch target genes, including *Hes1, Hey1* and *Hey2,* in Lin-cells (C), and HSC activation-associated genes, including *CDK6* and *MYC* in FL HSCs (LSK CD150^−^ CD48^+^) (D). Each dot represents one embryo (n=3).

**(E)** Bar graphs showing the frequency of LSK (left) and HSC (LSK CD48^−^ CD150^+^; right) contributions 16 weeks after primary, secondary and tertiary transplantations of E14.5 FL Lin-cells treated in vitro for 48 hours with DMSO (control) or compound E (CPDE) to inhibit Notch signaling. Each dot represents one mouse (n=3-8).

**(F)** Bars plots show the contribution to BM lineages including myeloid (Ly6C/Ly6G^+^ or Gr1^+^), erythroid (Ter119^+^), B (B220^+^ or CD45R^+^), and T (CD3^+^) cells, 16 weeks after primary (left), secondary (middle) and tertiary (right) transplantation of E14.5 FL Lin-cells treated in vitro for 48 hours with DMSO (control) or compound E (CPDE) to inhibit Notch signaling. Each dot represents one mouse (n=3-8).

**(G)** After 48 hours of Notch inhibition with CPDE in vitro, FL EPCR^+^ HSCs (LSK CD48^−^ CD150^+^ EPCR^+^) were sorted as single cells into individual wells and maintained in culture without further treatment. Experimental scheme (top). Single-cell (SiC) division assay showing manual quantification of cell number per well at 24 hours (left) and 48 hours (right). N = 2 independent experiments.

**(H)** Bar graphs showing the frequency of HSC (LSK CD48^−^ CD150^+^; left) and LSK (right) contributions 16 weeks after tertiary transplantations of adult BM cells treated in vitro for 48 hours with DMSO (control) or compound E (CPDE) to inhibit Notch signaling. Each dot represents one mouse (n=6).

**(I)** Bars plots show the contribution to BM lineages including myeloid (Ly6C/Ly6G^+^ or Gr1^+^), erythroid (Ter119^+^), B (B220^+^ or CD45R^+^), and T (CD3^+^) cells, 16 weeks after tertiary transplantation of adult BM Lin-cells treated in vitro for 48 hours with DMSO (control) or compound E (CPDE) to inhibit Notch signaling. Each dot represents one mouse (n=6).

## MATERIALS AND METHODS

### Mouse models

All mice used in this study were either of the C57BL/6 strain background or CD-1 mice. Animals were housed in groups of two to five per cage under a 12-hour light/dark cycle at a constant temperature of 23°C, humidity of 40-60% with ad libitum access to water and food (J. Rettenmaier & Söhne GmbH + Co KG; RM3 diet was provided to breeding pairs and mice up to nine weeks of age, after which they were switched to the RM1 maintenance diet).

All animal procedures were performed under specific pathogen-free (SPF) conditions, in compliance with the institutional and governmental guidelines for the care and use of laboratory animals, as established by the Animal Care Committee of the Generalitat de Catalunya. All experimental protocols were approved by the Committee for Animal Experimentation of the Barcelona Biomedical Research Park (PRBB).

Adult mice were euthanized by CO₂ inhalation, while pregnant females were euthanized by cervical dislocation. Postnatal mice up to postnatal day 9 (P9) were euthanized by decapitation, as their elevated levels of fetal hemoglobin make them less sensitive to CO₂ exposure.

For timed matings, females aged 2 to 6 months and males aged 2 to 12 months were used. The presence of a vaginal plug was designated as embryonic day 0.5 (E0.5). Embryos obtained from these matings were genotyped and subsequently used for downstream analyses.

Mice used were C57BL/6J (Jackson Laboratories, Strain #000664), CD-1 IGS (Charles Rivers, Strain #022), Vav-Cre (B6.Cg-Tg(VAV1-cre)1Graf/MdfJ) (Jackson Laboratories, Strain #035670), RBPj^flox^ (C57BL/6J-Rbpj^em2Lutzy^/J) (Jackson Laboratories, Strain # 034200), eEF1a^NICD^ (C57BL/6J-Tg(eEF1a-NICD,-loxP-stop-loxP)) mice were kindly provided by Dr. Iannis Aifantis, ROSA26^M2rtTa/M2rtTA^ (B6.Cg-Gt(ROSA)26Sor^tm1(rtTA*M2)Jae^/J) (Jackson Laboratories, Strain #006965), pTRE-H2BGFP^+/GFP^ (Tg(tetO-HIST1H2BJ/GFP)47Efu/J (Jackson Laboratories, Strain #005104).

### Genotyping PCR

An ear punch or a small fragment of embryonic tissue was collected and transferred into a PCR tube containing 100 μl of NaOH solution, then incubated at 98°C for 30 minutes to denature the DNA. The reaction was neutralized by adding 100ul of Tris-HCl buffer (pH of 7.4-8.0). After centrifugation at 10,000 rpm, 1 μl of the supernatant was used as a template for genotyping PCR. Primers used for genotyping were detailed in Table MM1.

**Table MM1.**
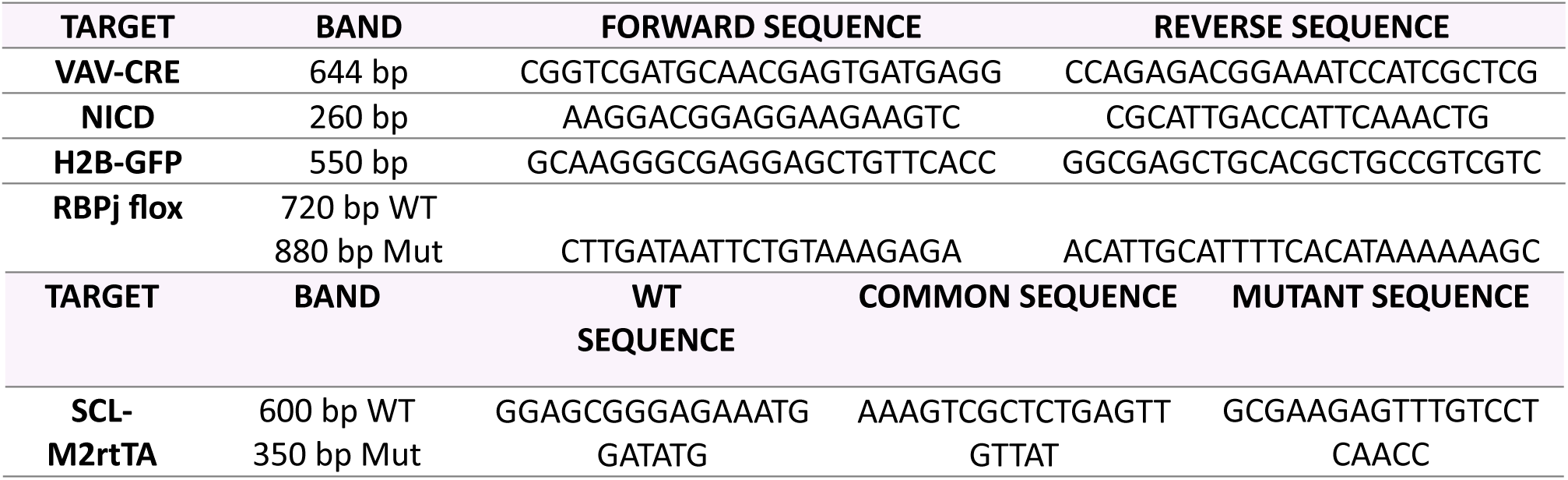
List of primers for mice genotyping.

### AGM, thymus, spleen, FL and BM analysis

AGM region from embryos were dissected in PBS and single-cell suspensions were generated by mechanical dissociation with a syringe and needle of 21G (BD Microlance, Cat #304432).

Fetal livers were collected from E12.5, E14.5, E16.5 embryos or newborns. Dissected livers were mechanically dissociated by gently pressing the tissue through a 40 μm cell strainer using the plunger of a 1 ml syringe into PBS with 10% fetal bovine serum (FBS). Spleen and thymus were processed in the same way as livers.

For BM isolation, femurs, tibias, hip bones and/or backbone were harvested, and marrow cells were obtained by crushing the bones in cold PBS supplemented with 10% FBS using a mortar and pestle. The resulting cell suspension was filtered through a 40 μm cell strainer to obtain a single-cell suspension.

### Lineage-positive cell depletion from FL and BM

Cells isolated from FL and BM were washed with PBS and incubated with a cocktail of biotin-conjugated antibodies targeting lineage markers (BD Biosciences, Cat# 559971) at a dilution of 1:200 for 20 minutes at 4°C. The lineage markers included Biotin Hamster anti-Mouse CD3e to identify T cells, biotin rat anti-mouse CD11b (or Mac1) for myeloid cells, biotin rat anti-mouse CD45R (or B220) for B cells, biotin rat anti-mouse Ly-6G and Ly-6C (or Gr1) for myeloid cells and neutrophils and biotin rat anti-mouse Ter119 for erythroid cells. Notably, CD11b was excluded from the lineage antibody cocktail during FL and newborn stages due to low CD11b expression levels on HSCs.

Following antibody labeling, cells were incubated with anti-biotin microbeads (Miltenyi Biotec, Cat#130-090-485) (1:100) and subjected to magnetic separation using LS columns (Miltenyi Biotec, Cat #130-042-401). The lineage-negative (Lin-) fraction, consisting of unlabeled cells, was collected for downstream applications.

### Peripheral blood analysis

Peripheral blood was collected via submandibular bleeding using heparinized capillary tubes (#22-362566, Fisherbrand), followed by red blood cell lysis with ACK lysing buffer (Lonza, Cat# BP10-548E).

The following antibodies were used for staining hematopoietic lineages from peripheral blood, BM or spleen: APC-Cy7 anti-CD45.1 (1:200), FITC anti-CD45.2 (1:200), PE anti-CD3 (1:400), PE-Cy7 anti-B220 (1:400), BV605 anti-Ter119 (1:200), APC anti-Gr1 (1:200) and APC anti-Mac1 (1:200). 1 μg/mL DAPI (1:200) (Biotium, Cat. No. BT-40043) was used as viability marker. The antibodies used are shown in Table MM2.

**Table MM2.** List of antibodies.

| ANTIBODY | COMPANY | CLONE | CATALOGUE<br>NUMBER |
| --- | --- | --- | --- |
| Biotin-conjugated CD3e | BD Biosciences |  | 51-01082J |
| Biotin-conjugated CD11b | BD Biosciences |  | 51-01712J |
| Biotin-conjugated CD45R | BD Biosciences |  | 51-01122J |
| Biotin-conjugated Ly-6G and Ly-6C | BD Biosciences |  | 51-01212J |
| Biotin-conjugated Ter-119 | BD Biosciences |  | 51-09082J |
| Biotin-conjugated Ter-119 | Invitrogen | TER-119 | 13-5921-82 |
| BrdU-APC | BD Pharmigen |  | 51-23619L |
| **CD11B / Mac1-APC** | BD Pharmingen | M1/70 | 553312 |
| **CD117 / c-kit-APC** | BD Pharmingen | 2B8 | 553356 |
| **CD117 / c-kit-APC-eFluor780** | eBioscience | 2B8 | 47-1172 |
| **CD117 / c-kit-FITC** | BD Pharmingen | 2B8 | 561680 |
| **CD135 / FLT3-Pe-Cy7** | BD Pharmingen | A2F10.1 | 567594 |
| **CD135 / FLT3-PE-CF594** | BD Biosciences | A2F10.1 | 562537 |
| **CD150 / SLAM-APC** | Biolegend | TC15-12F12.2 | 115909 |
| **CD150 / SLAM-PE-Cy5.5** | Biolegend | TC15-12F12.2 | 115912 |
| **CD150 / SLAM_BV785** | Biolegend | TC15-12F12.2 | 115937 |
| **CD201 (EPCR)-APCeFluor780** | eBioscience | eBio1560 | 47-2012-82 |
| **CD201 (EPCR)-PerCP-eFluor710** | eBioscience | eBio1560 | 46-2012-80 |
| **CD3-PE** | BD Biosciences | 145-2C11 | 553063 |
| **CD31-PE** | BD Pharmingen | MEC13.3 | 553373 |
| **CD34-PE-Cy7** | Biolegend | MEC14.7 | 119326 |
| **CD45-APC** | BD Pharmingen | 30-F11 | 559864 |
| **CD45.1-APC-eFluor 780** | eBioscience | A20 | 47-0453 |
| **CD45.2-FITC** | eBioscience | 104 | 11-0454 |
| **CD45.2-PerCP-Cy5.5** | BD Pharmingen | 104 | 552950 |
| **CD45R / B220-PE-Cy7** | BD Pharmingen | RA3-6B2 | 552772 |
| **CD48-Pacific Blue** | BioLegend | HM48-1 | 103418 |
| **CD48-PE** | BD Pharmingen | HM48.1 | 560731 |
| **CD48-PE-Cy7** | BD Pharmingen | HM48.1 | 560731 |
| **Gr1 / Ly-6g/Ly-6C** | Invitrogen | RB6-8C5 | 17-5931-82 |
| **KI67-Alexa Fluor 700** | BD Pharmingen | B56 | 561277 |
| **Sca-1(Ly-6Aa/E)-FITC** | BD Pharmingen | E13-161.7 | 553335 |
| **Sca-1(Ly-6Aa/E)-PE** | BD Pharmingen | D7 | 553108 |
| **Sca-1(Ly-6Aa/E)-PE-Cy7** | BD Pharmingen | D7 | 558162 |
| **Streptavidin-BV605** | Biolegend |  | 405229 |
| **Streptavidin-PE-CF594** | BD Biosciences |  | 562318 |
| **Streptavidin-PECP-Cy5.5** | Invitrogen |  | 2527222 |

### Competitive transplantation assays

C57BL/6 mice were used as recipients following two rounds of irradiation by a four-hour interval: male received a total dose of 8.5 Gy (4 Gy + 4.5 Gy), while females were irradiated with a total of 9 Gy (2 x 4.5 Gy). Mice were treated with the antibiotic enrofloxacin (10mg/kg) for 21 days post-transplantation.

In all experiments, age-matched animals of both sexes, between 8 and 12 weeks old, were employed. Varying numbers of donor cells were transplanted via retro-orbital injection, together with different quantities of competitor cells in primary transplantation assays (see below for detailed cell numbers per experiment). In secondary transplantations, BM cells were collected from primary recipients at 16 weeks post-transplantation, and a total of 1 million cells were transplanted into each irradiated secondary recipient.

Donor-derived peripheral blood chimerism was analyzed by flow cytometry at 4-, 8-, 12-, and 16-weeks post-transplantation. At 16 weeks, lineage analysis of cells isolated from BM, spleen, and peripheral blood was performed using flow cytometry with lineage-specific antibodies. Hematopoietic populations including Lin-, LSK and HSCs (LSK CD48^−^ CD150^+^) were analyzed from BM at 16 weeks post-transplantation using flow cytometry. The following antibodies were used: BV605-conjugated streptavidin, APC-Cy7 anti-c-Kit, PE anti-Sca1, APC anti-CD150, PE-Cy7 anti-CD48 and FITC anti-CD45.2. 1 μg/mL DAPI (1:200) (Biotium, Cat. No. BT-40043) was used as viability marker. Data acquisition was conducted on a BD Fortessa cytometer and analyzed using FlowJo software version 10. The antibodies used are shown in Table MM2.

#### RBPJ^f/f^VAV^CRE^ BM competitive transplantation assay

A total of 1×10^5^, 2×10^5^, 4×10^5^ and 7×10^5^ BM cells from RBPj^vav1^ WT and KO C57BL/6J (CD45.2) donor mice were co-transplanted with 2×10^5^ competitor BM cells (CD45.1) into lethally irradiated C57BL/6J (CD45.1) recipient mice. After 16 weeks post-transplantation, 1×10^6^ total BM cells from primary recipients were collected and transplanted into lethally irradiated secondary recipients.

#### RBPJ^f/f^VAV^CRE^ FL competitive transplantation assay

A total of 1×10^5^, 1×10^4^ and 1×10^3^ E14.5 FL cells from RBPj^vav1^ WT and KO C57BL/6J (CD45.2) donor mice were co-transplanted with 2×10^5^ competitor BM cells (CD45.1) into lethally irradiated C57BL/6J (CD45.1) recipient mice. Analysis was performed after 16 weeks of transplantation.

#### Competitive transplantation assay using Compound E-treated E14.5 FL or BM linage-negative cells

Lineage-negative (Lin⁻) cells were isolated from E14.5 FLs or adult BM and treated *in vivo* with Compound E or vehicle control (DMSO) for 48 hours. Following 48 hours of treatment, 5×10^4^ Lin-cells from FL or BM (CD45.2) were co-transplanted into lethally irradiated C57BL/6J (CD45.1) recipient mice along with 5×10^4^ total BM competitor cells (CD45.1). At 16 weeks post-transplantation, 1×10^6^ total BM cells from primary and secondary recipients were collected and transplanted into lethally irradiated secondary and tertiary recipients, respectively.

#### Competitive transplantation assay using postnatal day 5 (P5) H2B-GFP^HIGH^ and H2B-GFP^LOW^ cells from BM and liver

Following doxycycline administration at embryonic days E8.5, E9.5 and 10.5, a total of 29 HSCs (LSK CD48^−^ CD150^+^) were sorted from postnatal day 5 (P5) liver and BM based on H2B-GFP fluorescence intensity (high and low) from C57BL/6J ROSA26^M2rtTA/M2rtTA^;pTRE-H2BGFP^+/H2B-GFP^ mice (CD45.1/CD45.2). Sorted HSCs were co-transplanted with 5×10^4^ total BM competitor cells (CD45.2) into lethally irradiated C57BL/6J (CD45.2) recipient mice.

### Limiting Dilution Analysis (LDA) by Extreme Limiting Dilution Analysis (ELDA)

Limiting dilution analysis (LDA) was performed to estimate the frequency of HSCs in competitive transplantation assays. This method enables the calculation of the Competitive Repopulating Unit (CRU), defined as the number of functional HSCs within a sample that are capable of long-term hematopoietic reconstitution in recipient mice following transplantation. LDA is based on the single-hit Poisson model, which assumes that the engraftment of a single functional HSC is sufficient to reconstitute hematopoiesis and produce a detectable level of donor-derived differentiated cells above a defined threshold in recipient animals. The analysis was conducted using the ELDA (Extreme Limiting Dilution Analysis) software(Hu and Smyth 2009), based on the number of transplanted cells (dose), the number of recipients per dose and the number of successful engraftments (positive response). Engraftment levels below 10% were considered as negative and excluded from the frequency calculation.

### *In vivo* BrdU administration and expression analysis

Bromodeoxyuridine (BrdU) was administered to mice via intraperitoneal injection at a dose of 1.5 mg per mice, using a 10 mg/ml BrdU solution in sterile PBS with a final volume of 200 μl, to performed long-term label-retention cell assay *in vivo*. A total of 100 μl diluted BrdU was injected intraperitoneally on each side of the abdomen. The injection was performed at three consecutive embryonic days E9.5, 10.5 and E11.5. At defined time points post-injection, embryos or postnatal mice were collected and processed for downstream analyses of BrdU expression by flow cytometry.

Fetal or postnatal livers were harvested and dissociated into single-cell suspensions. Lin-cells were isolated by lineage depletion and subsequently stained to identify hematopoietic subpopulations, including Lin-, LSK, HSCs (LSK CD48^−^ CD150^+^) and EPCR^+^ HSCs. The following antibodies were used for surface staining: PE-Texas Red-conjugated streptavidin (1:400), FITC anti-c-Kit (1:200), PE anti-Sca1 (1:400), PE-Cy5.5 anti-CD150 (1:400), PE-Cy7 anti-CD48 (1:400) and APC-Cy7 anti-EPCR (1:200). Following this, cells were fixed and permeabilized using the BrdU Flow Kit (BD Pharmigen, Cat #552598) according to the manufacturer’s instructions. DNA was denatured by treatment with DNase for 1 hour at 37°C to allow detection of incorporated BrdU. Cells were then incubated with APC anti-BrdU antibody for 30 minutes at room temperature. For DNA content analysis, cells were stained overnight at 4°C with DAPI (5 μg/ml; 1:1000 dilution; Biotium, Cat. #BT-40043). Flow cytometry analysis was performed using BD LSR Fortessa Flow Cytometer (BD Bioscience). The antibodies used are shown in Table MM2.

### Doxycycline delivery *in vivo* and H2B-GFP expression analysis

Timed pregnancies were established by crossing pTRE-H2BGFP^+/GFP^ males with ROSA26^M2rtTa/M2rtTA^ females. Pregnant females exhibiting a vaginal plug were selected for treatment. Doxycycline (DOX) (Tocris Bioscience, Cat #4090) was administered via intraperitoneal injection on three consecutive embryonic days (E8.5, E9.5, and E10.5) to label cells. An injection of DOX at 1 mg/ml per 20 g of body weight, diluted in PBS to a final volume of 200 μl, was administered. A total of 100 μl diluted DOX was injected intraperitoneally on each side of the abdomen. At defined time points post-injection, embryos or postnatal mice were collected and processed for downstream analysis of long-term label-retaining cells by quantifying H2B-GFP expression using spectral flow cytometry.

To identify hematopoietic subpopulations, including lineage-negative (Lin-) cells (defined by CD3^+^, B220^+^, Gr1^+^, Ter119^+^), LSK cells (c-Kit^+^ Sca1^+^), HSCs (LSK CD34^−^ CD135^−^ CD48^−^ CD150^+^), EPCR^+^ HSCs, MPP1 (LSK CD34^+^ CD135^−^ CD48^−^ CD150^+^), MPP2 (LSK CD34^+^ CD135^−^ CD48^+^ CD150^+^), MPP3 (LSK CD34^+^ CD135^−^ CD48^+^ CD150) and MPP4 (LSK CD34^+^ CD135^+^ CD48^+^ CD150^−^), the following antibodies were used for surface staining: BV605-conjugated streptavidin (1:400), APC anti-c-Kit (1:200), PE anti-Sca1 (1:400), BV785 anti-CD150 (1:200), Pacific Blue anti-CD48 (1:200), PE-Cy7 anti-CD34, PE-CF594 anti-CD135 and APC-Cy7 anti-EPCR (1:200). Staining was performed for 20 minutes at 4°C. The antibodies used are shown in Table MM2.

Due to the high fluorescence intensity of the H2B-GFP reporter, all laser powers had to be reduced 90-fold during spectral flow cytometry acquisition to avoid signal saturation and ensure accurate detection.

### *In vitro* culture treatments

For *in vitro* cell treatments, the following reagents were used: 1μM Compound E or γ-Secretase Inhibitor XXI (Sigma-Aldrich Cat #565790), 10μM or 50μM DAPT or γ-Secretase Inhibitor IX (Sigma-Aldrich Cat #565770), 10 µg/ml Notch1 hIgG1 blocking antibody (Custom reagent provided by Genentech Batch/lot #PUR567908), 10 µg/ml Notch2 mIgG2 blocking antibody (Custom reagent provided by Genentech Batch/lot #PUR611799), 10 µg/ml IgG from human serum (hIgG) (Sigma-Aldrich Cat #I4506), 10 µg/ml IgG from mouse serum (mIgG) (Sigma-Aldrich Cat #I8765) and Dimethyl sulfoxide (DMSO) (Sigma-Aldrich Cat #34869).

### Lineage-negative populations culture

E14.5 FL or adult BM were processed and red blood cells were lysed using ACK lysing buffer (Lonza, Cat# BP10-548E). Lin-cells were obtained and seeded into 6-well plates at a density of 0.6×10^6^ cells per well and 2 mL of culture medium per well. The culture medium consisted of StemSpan™ SFEM medium (Stem Cell Technologies Cat# 09600) supplemented with 2 mM L-glutamine (Biological Industries Cat #03-020-1A), 100 U/mL penicillin and streptomycin (Biological Industries Cat #03-031-1B), 30 ng/ml mouse Flt3 ligand (Peprotech Cat# 250-31L-10UG), 50 ng/ml mouse SCF (Peprotech Cat# 250-03-10UG) and 25 ng/ml mouse thrombopoietin (TPO) (Peprotech Cat# 315-14-10UG). Culture medium was refreshed every 3 days.

### Single cell division assay

Hematopoietic cells were isolated from E14.5 FL, and lineage-positive cells were depleted. The resulting FL Lin-cells were culture in vitro for 48 hours in the presence of 1 μM Compound E (gamma-secretase inhibitor) or DMSO (control), following the same culture condition previously detailed. After transient Notch inhibition, Lin-cells were collected and sorted in single cells (LSK CD48^−^ CD150^+^ EPCR^+^) into 96-well plates containing StemSpan medium supplemented with cytokines, as previously described. Cells were maintained in culture without any additional treatment, to mimic transplantation settings, and the number of cells per well was manually counted at 24- and 48-hours post-sorting.

### Cell cycle analysis

Hematopoietic cells were harvested from BM and FL and stained for lineage-positive populations using a cocktail of biotin-conjugated mouse lineage antibodies (BD Pharmingen, Cat. No. 559971) (1:200). The lineage markers used were CD3, B220, Mac-1, Gr-1, and Ter119 for BM cells, and CD3, B220, Gr-1, and Ter119 for FL cells (excluding Mac-1) for 20 minutes at 4°C. Following this, hematopoietic cells including Lin-, LSK and HSCs (LSK CD48^−^ CD150^+^) were stained with BV605-conjugated streptavidin (BioLegend Cat #405229) (1:200), APCCy7 anti-c-Kit (BioLegend Cat #135136) (1:200), PE-Cy7 anti-Sca1 (BioLegend Cat #108114) (1:400), APC anti-CD150 (BioLegend Cat #115910) (1:200), PE anti-CD48 (BS Biosciences Cat# 552855) (1:400) antibodies for 20 minutes at 4°C.

Cells were then fixed and permeabilized using the FIX & PERM Cell permeabilization kit (Invitrogen, Cat #GAS004) following manufacturer’s instructions. Intracellular staining was performed with Alexa Fluor 700 (AF700) anti-Ki-67 antibody (1:100) for 20 minutes at room temperature. Finally, cells were incubated overnight at 4°C with DAPI (5 μg/ml; 1:1000 dilution) (Biotium, Cat #BT-40043) for DNA content assessment. Flow cytometry analysis was performed using BD LSR Fortessa Flow Cytometer (BD Bioscience). The antibodies used are shown in Table MM2.

### Flow cytometry analysis and cell sorting

Flow cytometry analyses were conducted on either a BD LSRFortessa Cell Analyzer or a BD LSR II Flow Cytometer (BD Biosciences).

Cells were purified by fluorescence-activated cell sorting (FACS) using a BD FACSAria II Cell Sorter (BD Biosciences). Sorting was performed at approximately 4,000 events per second using an 85 μm nozzle. Hematopoietic cells from FL or BM were stained with the cocktail of biotin-conjugated antibodies targeting lineage markers (BD Biosciences, Cat# 559971) at a dilution of 1:200 for 20 minutes at 4°C as explained before. After that, the following FACS antibodies were used: BV605-conjugated streptavidin (BioLegend Cat #405229) (1:200), FITC anti-c-Kit (BD Pharmigen Cat #561680) (1:200), PE anti-Sca1 (BD Pharmigen Cat #553335) (1:400), APC anti-CD150 (BioLegend Cat #115910) (1:200), PE-Cy7 anti-CD48 (BD Pharmigen Cat# 560731) (1:400) and APC-eFluor780 anti-EPCR (eBioscience Cat#47-2012-82) antibodies for 20 minutes at 4°C. Viability staining was performed using 5 μg/mL DAPI (Biotium, Cat. No. BT-40043). The antibodies used are shown in Table MM2.

Data acquisition and analysis were performed using Diva software version 6.1.2 (BD Biosciences) and FlowJo software version 10.0.6 (Tree Star).

### Spectral flow cytometry analysis and cell sorting

Spectral flow cytometry analysis was conducted using a Cytek Aurora Spectral Flow Cytometer (Cytek Biosciences). Spectral flow cytometry sorting was conducted using a Cytek Aurora CS 5L Spectral Flow Cytometer (Cytek Biosciences). Sorting was performed at approximately 4,000 events per second using a 70 or 85 μm nozzle.

Prior to data acquisition, spectral unmixing was performed using single-stained compensation controls to accurately separate fluorochrome emissions. Viability staining was performed using 1 μg/mL DAPI (1:200) (Biotium, Cat. No. BT-40043). Data acquisition and spectral unmixing were performed using SpectroFlo software (Cytek Biosciences). Further data analysis was conducted using FlowJo software version 10.0.6 (Tree Star).

### RNA isolation, cDNA synthesis and quantitative RT-PCR

Total RNA was extracted from cells using the RNeasy Plus Mini Kit (Qiagen, Cat. No. 74136) or the RNeasy Micro Kit (Qiagen, Cat. No. 74004), according to the manufacturer’s instructions. RNA concentration was measured using a NanoDrop spectrophotometer (Thermo Fisher Scientific, Cat. No. ND2000CLAPTOP). A total of 2 μg of RNA was reverse transcribed into cDNA using the Transcriptor First Strand cDNA Synthesis Kit (Roche, Cat. No. 04897030001), following the manufacturer’s protocol.

Quantitative real-time PCR (qRT-PCR) was performed in technical triplicates using the SYBR Green I Master Kit (Roche, Cat. No. 04887352001) on the LightCycler 480 System (Roche). Gene expression levels were calculated using the 2^−ΔCT^ method, normalized to the average CT of the Tbp or Gapdh housekeeping genes. Primer sequences are listed in Table MM3.

**Table MM3.**
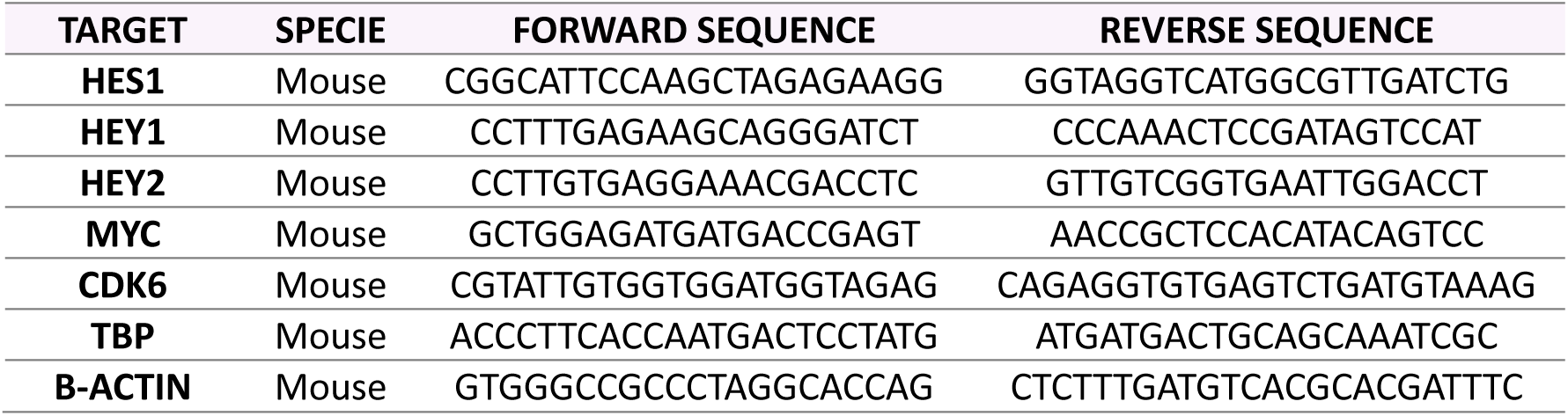
List of primers for RTqPCR analysis.

### cDNA preamplification

In the case of low-input cDNA obtained from fewer than 500 cells, SsoAdvanced™ PreAmp Supermix (BioRad Cat#1725160) was used to preamplify the cDNA following the manufacturer’s instructions. Briefly, 5 ng of cDNA were mixed with target-specific primers at a final concentration of 500nM and 1x SsoAdvanced PreAmp Supermix in a final volume of 20 μl. Amplification was performed in a PCR thermocycler under the following conditions: 95 °C for 3 min, followed by 12–14 cycles of 95 °C for 15 s and 60 °C for 4 min. The preamplified cDNA was diluted 1:5 in nuclease-free water and used subsequently used for qPCR analysis.

### Bulk RNA sequencing samples

Three different datasets were considered for bulk RNA-seq:

#### (i)#Adult BM HSCs isolated from RBPj^vav1^ KO and WT mice

Adult BM was processed from RBPj^vav1^ WT and KO mice, as previously described. Each mouse was analyzed individually and 300 HSCs (LSK CD48^−^ CD150^+^) were isolated by FACS. A total of 3 to 4 biological replicates were analyzed.

#### (ii)#E14.5 FL HSCs isolated from NICD^vav1^ and WT embryos

E14.5 FL were processed from embryos with constitutive activation of Notch (NICD^VAV-CRE^) and from WT embryos, as previously described. Each embryo was analyzed individually and 300 HSCs (LSK CD48^−^ CD150^+^) were isolated by FACS. Only male embryos were included in the study. A total of 2 to 3 biological replicates were analyzed for RNA-seq.

#### (iii)#P5 neonatal liver and BM HSCs isolated from high or negative H2B-GFP levels

P5 neonatal liver and BM were processed from H2B-GFP;M2rtTA mice with high or negative GFP intensity, as previously described. Each mouse was analyzed individually and 300 LT-HSCs (LSK CD48^−^ CD150^+^ CD34^−^ CD135^−^) were isolated by FACS. A total of 3 biological replicates were analyzed for RNA-seq.

### Bulk RNA sequencing analysis

#### RNA EXTRACTION AND LIBRARIES PREPARATION

Total RNA from three to four independent samples was extracted using the RNeasy Plus Mini Kit (Qiagen; Cat #180492) following manufacturer’s instructions and for library pre-preparation for (i) RBPj^vav1^ WT and KO BM HSCs. In the case of (ii) WT and NICD^VAV-CRE^ E14.5 FL HSCs and (iii) P5 H2B-GFP high and negative liver and BM HSCs, libraries were prepared using QIAseq Ultralow Input Library Kit (Qiagen, Cat #180492). In all cases, RNA concentration and integrity were determined using Agilent Bioanalyzer.

Libraries were prepared according to standard protocols. The Clontech SMARTer kit for low input material was used for both FL datasets. The resulting cDNA libraries were sequenced on different Illumina platforms depending on the dataset. Specifically: (i) RBPj^vav1^ BM dataset, sequencing depth ranged between 35M and 59M reads (average 47M reads) per sample (50bp single-end reads) in platform Illumina HiSeq 2500. (ii) NICD FL dataset sequencing depth ranged between 27M and 67M reads (average 35M reads) per sample (50bp paired-end reads) in platform Illumina HiSeq 2500. And finally, (iii) P5 H2B-GFP high and negative liver and BM HSCs dataset sequencing depth ranged between 51M and 68M reads (average 62M reads) per sample (150bp paired-end reads) in platform NextSeq6000.

#### DATA PRE-PROCESSING

In case of NICD FL and RBPj^vav1^ BM datasets, quality control was performed on raw data with the FASTQC tool (v0.11.9). Raw reads were trimmed to remove adapter presence with Trimgalore (v0.6.6) (The Babraham Institute by @FelixKrueger). Default parameters were used except for a minimum quality of 15 (Phred score) and an adapter removal stringency of 3bp overlap. Trimmed reads were aligned to reference the genome with STAR aligner tool (v2.7.8)(Dobin, Davis et al. 2013). STAR was executed with default parameters except for the number of allowed mismatches which was set to 1. The required genome index was built with corresponding GRCm38 gtf and fasta files retrieved from Ensembl (http://ftp.ensembl.org/pub/release-102/). Obtained BAM files with uniquely mapped reads were considered for further analysis except for one wild-type sample (labeled as S17) from the NICD FL dataset which corresponding BAM file was subsampled to its 40% using samtools (v1.15)(Danecek, Bonfield et al. 2021). Raw gene expression was quantified using featureCounts tool from subRead software (v2.0.1) with exon as feature(Liao, Smyth et al. 2014). In the case of Compound E FL dataset, all steps were the same except for the trimming and additional parameters during alignment. For trimming, cutadapt (v4.2) tool was used to remove Illumina universal adapters (3’ end) and the Clontech SMARTer IIA oligo (AAGCAGTGGTATCAACGCAGAGTAC) presence at 5’ end (Martin 2011), Regarding alignment, following parameters were included: --peOverlapNbasesMin 5, --peOverlapMMp 0.01, -- outFilterMismatchNoverLmax 0.01, --outFilterScoreMinOverLread 0.5 and -- outFilterMatchNminOverLread 0.5.

#### DIFFERENTIAL EXPRESSION ANALYSIS

For all three datasets, the raw counts matrix (including 55,487 genes) was imported into the R Statistical Software environment (v4.2.1) for statistical analysis. Prior to statistical analysis, those genes with less than 5 raw counts across all samples under test (per dataset) were removed. For visualization purposes, counts were normalized by the variance-stabilizing transformation method as implemented in DESeq2 R package (v1.38.3)(Love, Huber et al. 2014). Differential expression analysis (DEA) was conducted with DESeq2. Obtained log2 fold change values were shrunken with apeglm shrinkage estimator R package (v1.20.0)(Zhu, Ibrahim et al. 2019). Raw p-values were adjusted for multiple testing using the Benjamini-Hochberg False Discovery Rate (FDR)(Benjamini and Hochberg 2018). Differentially Expressed Genes (DEGs) for comparison of interest were called with adjusted p-values (FDR) < 0.05. Following the specific details per dataset are described: (i) in case of the RBPj^vav1^ WT and KO BM dataset, a total of 7 samples were available for analysis (3xKO and 4xWT). A total of 28,663 genes were kept after filtering very low expressed genes. Fitted statistical model included mouse gender and sample condition as covariable with RBPj^vav1^ WT as the reference. (ii) In the case of the Notch WT and overexpression of NICD FL HSCs dataset, a total of 5 samples were available for analysis (3xNICD and 2xWT). A total of 21,266 genes were kept after filtering very low expressed genes. Fitted statistical model included sample condition as covariable with WT as the reference. And finally, (iii) for P5 H2B-GFP high and negative liver and BM HSCs dataset, a total of 6 samples were available for analysis (3xGFP-negative 24h and 3xGFP-high). A total of 25,155 genes were kept after filtering very low expressed genes.

#### FUNCTIONAL ANALYSIS

Gene Set Enrichment Analysis (GSEA) was performed for comparisons of interest in each of the bulk RNA-seq datasets. We considered a total of 26 gene signatures previously associated to HSC dormancy, activated HSC/MPP, metabolic activity and cell cycle. Specifically they were retrieved from Table S1 (mouse) from Ishida et al.(Ishida, Mercoli et al. 2025) plus four gene sets from Wikipathways database(Agrawal, Balci et al. 2024) (WP234 GPCRs, WP374 Prostaglandin, WP157 Glycolysis and WP295 ETC) and one from Reactome database (R-MMU-72766 Translation) that were also highlighted in Cabezas-Wallscheid et al.(Cabezas-Wallscheid, Buettner et al. 2017). Additionally, Notch signaling Pathway from MSigDB HALLMARK(Castanza, Recla et al. 2023) and a custom extended version (adding Dll4, Notch4, Jag2, Hes5, Hes7, Hif1a, Nrarp, Dtx3, Nfkb2, Cdkn1a, Gata3 and Ptcra genes) was also considered for testing. The complete list of genes under test (28,663 genes in case of RBPj^vav1^ KO BM, 21,266 genes for NICD FL and 25,155 genes from H2B-GFP HSCs datasets) was ranked based on the shrunken log2 Fold Change obtained from DEA. GSEA was conducted through the fgseaMultilevel function from fgsea R package (v1.24.0) with default parameters (Korotkevich, Sukhov et al. 2021). A significance level of α=0.1.

### Single cell RNA sequencing data analysis

#### SAMPLES

FLs from embryonic day E14.5 were processed from RBPj^vav1^ WT and KO embryos, as described in Methods 3 and 4. Each embryo was analyzed individually and 50,000 to 80,000 FL LSK cells were isolated by FACS, following the protocol detailed in Method 16. Single-cell RNA sequencing 3’ (scRNA-seq) was performed using the Chromium Controller from 10x Genomics according to the manufacturer’s instructions.

#### DATA PREPROCESSING

10x sequencing raw reads were demultiplexed and aligned using 10x Genomics CellRanger v7.2.0 (cellranger count) under the default parameters which include intronic reads. Samples were aligned against mouse pre-built refdata-gex-mm10-2020-A reference transcriptome provided by 10x Genomics. Features, barcodes and expression matrices were obtained independently for each sample (1xRBPj^vav1^ WT and 1xRBPj KO). The RBPj^vav1^ WT sample was sequenced twice because of its lower sequencing depth compared to the RBPj KO sample. The average of recovered cells was 12,791 and 4,656 cells, the median genes per cell was 3,745 and 4,124 genes and the mean reads per cell was 32,162 and 41,761 reads, respectively for RBPj^vav1^ WT and KO samples.

#### QUALITY CONTROL

Generated filtered matrices were imported and merged into R (v4.3.3)(Team 2021) using Seurat package (v5.2.1)(Hao, Stuart et al. 2024). A Seurat object with a total of 32,285 genes and 17,447 cells was available prior to quality control (QC). Doublets were separately detected by sample with scDblFinder R package (v1.16.0)(Germain, Lun et al. 2021). For this purpose, random and cluster-based approaches, implemented in the same package, were used. Additionally, the latter was considered with the clusters identified by CellRanger software (graph-based). Those cells detected as doublets by two out of the three cases were finally labeled as doublets and removed from the dataset. For RBPj^vav1^ WT and KO conditions respectively, 8.7% and 6.7% of cells were removed. Next, cells were filtered based on the number of genes. Cells with fewer than less than 500 or more than 10k genes were excluded. Cells with more than 8% of mitochondrial gene content or with less than 2k counts were also discarded. Genes do not present in at least 5 cells were filtered out. Ribosomal genes were also discarded. Following quality control, a Seurat object with a total of 19,276 genes and 11,389 cells was used for downstream analysis.

#### DIMENSIONALITY REDUCTION

Samples were normalized using the SCTransform with the method glmGamPoi (package v1.14.3)(Ahlmann-Eltze and Huber 2021), no regressing any variable out. No sample integration was required. For dimensionality reduction and visualization, Uniform Manifold Approximation (UMAP)(Armstrong, Martino et al. 2021) was used. The runPCA and runUMAP functions were executed considering 35 principal components which corresponded to a cumulative captured data variance > 85%. PHATE(Moon, van Dijk et al. 2019) dimensionality reduction method (phateR_package v1.0.7) was also used for visualization purposes. Cell phase score was computed with the CellCycleScoring function and available S and G2M genes in the Seurat package. Expression values were imputed and smoothed with the MAGIC algorithm through Rmagic R package (v2.0.3.999)(van Dijk, Sharma et al. 2018).

#### CLUSTERING AND CELL TYPE ANNOTATION

Cell types were annotated by integrating our dataset with an already annotated mouse E14.5 FL dataset(Gao, Shi et al. 2022). Integration was conducted with harmony R package (v1.2.3)(Korsunsky, Millard et al. 2019). The integrated dataset contained 19,793 cells in total. To transfer the reference cell type labels to our dataset, we clustered integrated cells with different resolutions with the aim of obtaining clusters containing > 75% reference cells of the same type. For this purpose, FindNeighbors (35 dimensions) and FindClusters functions were used with default parameters except for the resolution. An initial resolution of 0.2 was conducted, obtaining 8 clusters with a percentage of common reference cells greater than 75%. These clusters were identified as CLP, Macrophages, Megakaryocytes, MegE progenitors and Neutrophils (clusters containing Endothelial cells and Hepatoblast were also annotated but corresponded exclusively to the reference dataset). Clusters remaining unannotated were further subdivided using the FindSubCluster function at a resolution of 0.25, resulting in 12 additional clusters. Applying the same criteria, 7 of these were assigned to specific identities, including CMP, Erythroblasts, Erythroid cells and HSC/MPPs. Clusters that were not annotated due to cell type heterogeneity were manually annotated by combining the two most prevalent reference cell types.

Manual validation of previous cell annotations was conducted by exploring the expression of marker genes already used in the reference dataset, including : HSC/MPP (Procr, Mecom, Hoxa9, Mycn, Hlf), Neutrophil (Gstm1, Elane, Ctsg, Fcnb), Macrophage1 (Cd68, Ly86, Fcgr1, Csf1r, Hmox1), Macrophage2 (Adgre1, C1qa), MegE progenitor (Gata1, Epor, Klf1, Car1), Erythroid cell (Hbb-bs, Hbb-bt, Hba-a2, Ermap, Alad), Mk (Vwf, Itga2b, Itgb3, Plek, Pf4), CMP (Mpo, Spi1, Cd34, Prtn3, Ccl9), CLP (Ccr9, Cd7, Notch1, Satb1, Cd79a), Basophil (Cpa3, Rnase12, Il6, Stx3, Alox15), and B cell (Blnk, Pax5, Ebf1, Vpreb1). As a result of this manual validation, two clusters previously identified as MegE progenitor and Megakaryocyte were re-annotated as Basophils.

Cells annotated as HSC/MPP were subsetted and clustered with 0.1 resolution obtaining two clusters. Each cluster was labelled as HSCs or MPPs based on the expression of known HSC markers related to dormancy (Hlf, Mecom, Procr, Mllt3, Meg3 and H19 as the *Dormancy* module) and activity (Cdk4, Cdk6, Myc, Mki67, Cd48, Cdk1, Top2a, Cdc20 and Cd34 as the *Activity* module), respectively. Scikit-learn trained hscScore model (v0.21.2)(Hamey and Gottgens 2019) was employed to infer stemness potential in HSCs.

The processes of normalization, dimensionality reduction and imputation of expression values were independently re-applied to each subset of the original dataset.

#### HSC CLUSTERING

Hierarchical clustering analysis was performed to classify HSCs into dormant HSCs and active HSCs (method Complete, distance metric Euclidean). Gene expression levels of HSC markers related to dormancy and activity, also used for HSC/MPP cell type annotation, were considered. Four main clusters were identified. Those exhibiting the highest expression of dormant markers and the lowest expression of active markers were annotated as dormant, while those with the opposite pattern were annotated as active. Clusters with a less defined expression pattern were annotated as intermediate.

#### GENE SIGNATURE SCORING AND VISUALIZATION

Data visualization was performed with the ggplot2 (v3.5.2) (Wickham 2016) ComplexHeatmap (v2.18.0)(Gu, Eils et al. 2016) and tidyplots (v0.2.2)(Engler 2025) R packages. A selected list of gene signatures were scored, specifically: adult HSC dormancy(Cabezas-Wallscheid, Buettner et al. 2017), Activated HSC/MPPs(Schonberger, Obier et al. 2022), serial-engrafting FL HSCs(Ishida, Mercoli et al. 2025), diapause(Boroviak, Loos et al. 2015, Duy, Li et al. 2021), low-output(Rodriguez-Fraticelli, Weinreb et al. 2020), high-output(Rodriguez-Fraticelli, Weinreb et al. 2020) and MYC-targets v1 and v2 from MSigDB HALLMARK(Castanza, Recla et al. 2023) database. Gene signature scores were calculated using the AddModuleScore function from the Seurat package over the MAGIC-imputed data.

#### STATISTICAL ANALYSIS

Statistical analysis was performed using R software. The Wilcoxon Test was applied to compare the expression of different signatures between groups. The statistical significance of each result is denoted as follows: ns for p > 0.05, * for p ≤ 0.05, ** for p ≤ 0.01, *** for p ≤ 0.001, and **** for p ≤ 0.0001.

### Division rate estimation of H2B-GFP-retaining HSCs

To estimate the number of cell divisions that HSCs from M2rtTA;H2B-GFP mice undergone during fetal and early postnatal development, we applied a GFP dilution–based mathematical model(Ganuza, Hall et al. 2022). Upon administration of a doxycycline (DOX) pulse in the H2B-GFP system, the fusion protein H2B-GFP is stably incorporated into chromatin. As labeled cells subsequently divide, the H2B-GFP content is equally segregated between daughter cells, leading to a progressive dilution of the GFP signal. This dilution results in a halving of the median fluorescence intensity (MFI) of GFP protein with each cell division. Thus, the division rate can be estimated using the formula N = log_2_ (Y_0_/Y_N_), where N = number of cell divisions, y_0_ = initial H2B-GFP MFI, and y_N_ = H2B-GFP MFI after N divisions. It was considered that H2B-GFP labeling remains active for approximately 24 hours following DOX withdrawal, thus H2B-GFP labeling will continues until E11.5 with the current treatment protocol.

Because progressive GFP dilution limits the accuracy of division estimates after multiple cell cycles, we further inferred the actual number of cell divisions (Nₐ) using a previously described correction model(Ganuza, Hall et al. 2022), in which Nₐ is calculated as an exponential function of the estimated value: Nₐ = e^(−0.757+0.541*N)^. This approach enables more accurate quantification of cumulative cell divisions within the dynamic range of the H2B-GFP system.

### Statistical analysis

Statistical analyses were performed using GraphPad Prism software (v9.0.0, Dotmatics) and p-value<0.05 is considered significant (****p-value<0.0001, ***pvalue<0.001, **p-value<0-01 and n.s. p-value > 0.05). A 95% confidence interval was used to define statistical significance, and significance level of α=0.05. Statistical parameters, including the standard deviations and statistical significance values, are provided in the corresponding figures and figure legends.

For comparisons between two groups, data normality was assessed using the Shapiro–Wilk test. Student’s t-test was applied when data followed a normal distribution, whereas the Wilcoxon rank-sum test was used for non-normally distributed data. For multiple group comparisons, either one-way ANOVA followed by Tukey’s post hoc correction or two-way ANOVA followed by Sidak’s multiple comparisons test was used, depending on the experimental design. In the case of categorical variables or proportional data involving small sample sizes, statistical significance was assessed using Fisher’s exact test.

