## Supplementary figures 1-6 for "Developmental programming of hematopoietic stem cell dormancy by evasion of Notch signaling"

### FIGURE S1

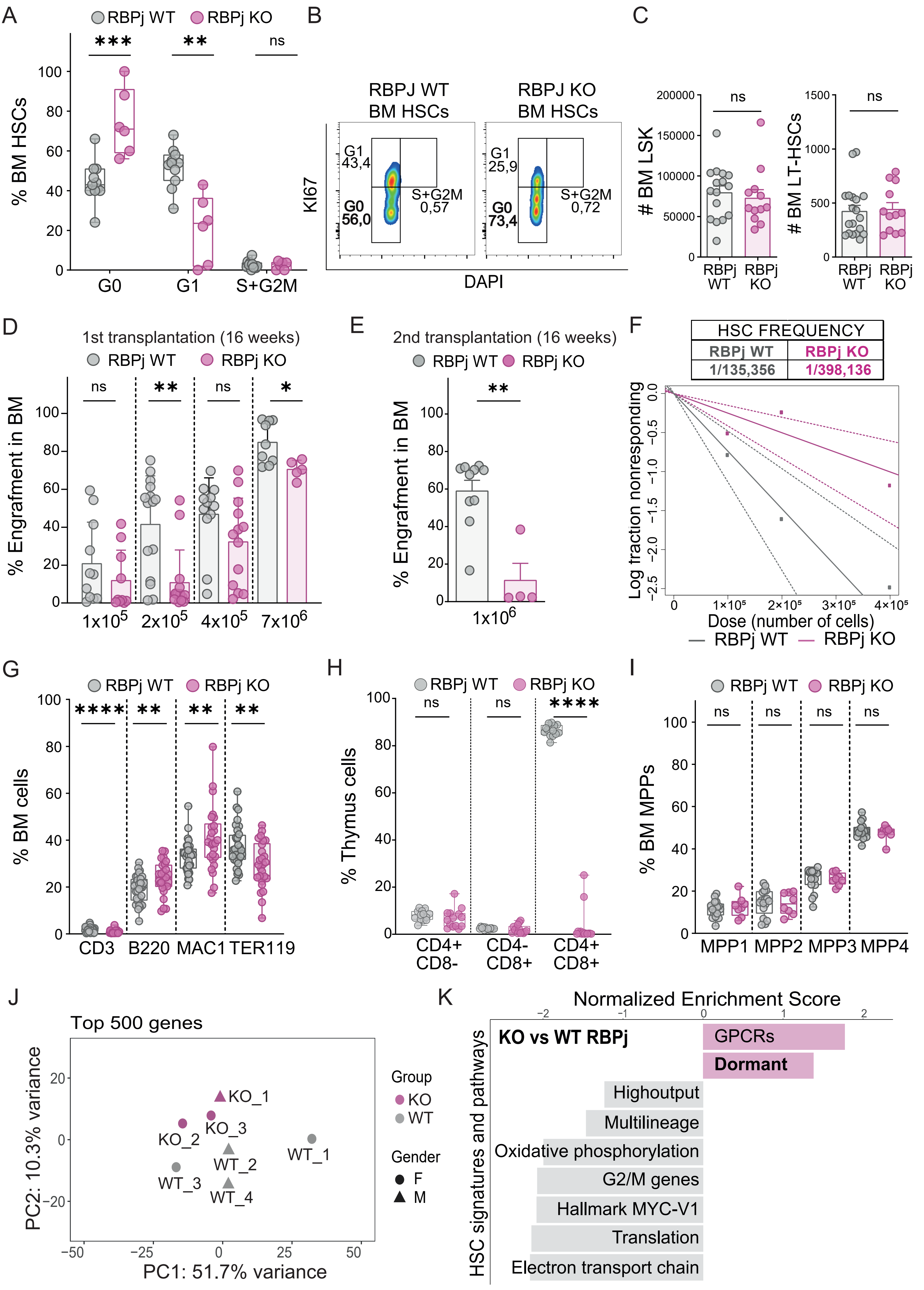

A
RBPj WT
RBPj KO
***
**
ns
% BM HSCs
G0
G1
S+G2M
B
RBPJ WT
BM HSCs
RBPJ KO
BM HSCs
KI67
G1
43,4
G0
56,0
S+G2M
0,57
G1
25,9
G0
73,4
S+G2M
0,72
DAPI
C
### BM LSK
### BM LT-HSCs
RBPj WT
RBPj KO
D
1st transplantation (16 weeks)
% Engrafment in BM
1x10^5
2x10^5
4x10^5
7x10^6
E
2nd transplantation (16 weeks)
1x10^6
F
HSC FREQUENCY
RBPj WT
RBPj KO
1/135,356
1/398,136
Log fraction nonresponding
Dose (number of cells)
G
% BM cells
CD3
B220
MAC1
TER119
****
H
% Thymus cells
CD4+
CD8-
CD4-
CD8+
CD4+
CD8+
I
% BM MPPs
MPP1
MPP2
MPP3
MPP4
J
Top 500 genes
PC2: 10.3% variance
PC1: 51.7% variance
KO_1
KO_2
KO_3
WT_1
WT_2
WT_3
WT_4
Group
KO
WT
Gender
F
M
K
Normalized Enrichment Score
HSC signatures and pathways
KO vs WT RBPj
GPCRs
Dormant
Highoutput
Multilineage
Oxidative phosphorylation
G2/M genes
Hallmark MYC-V1
Translation
Electron transport chain

### FIGURE S2

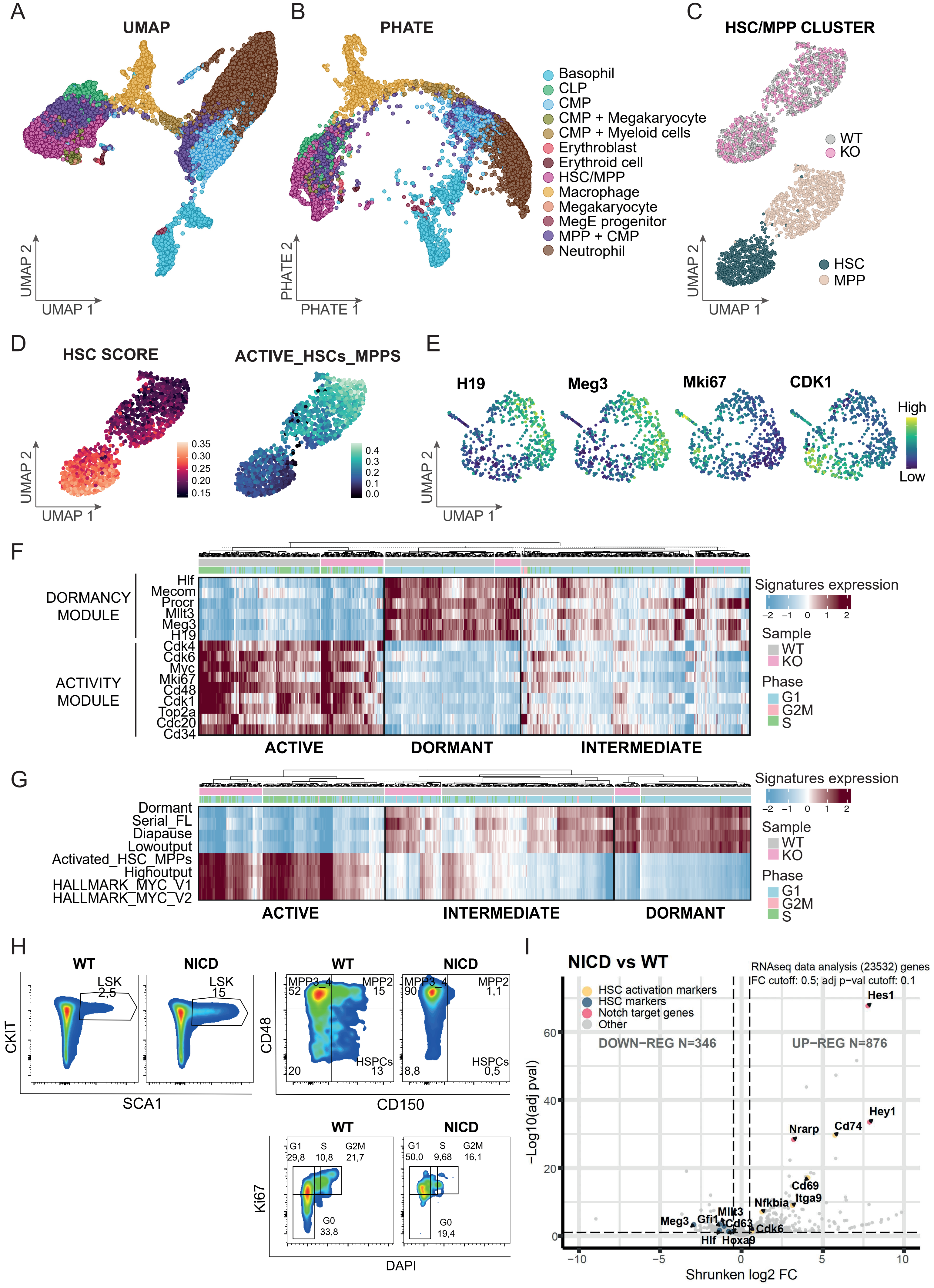
A
UMAP
UMAP 1
UMAP 2
B
PHATE
PHATE 1
PHATE 2
Basophil
CLP
CMP
CMP + Megakaryocyte
CMP + Myeloid cells
Erythroblast
Erythroid cell
HSC/MPP
Macrophage
Megakaryocyte
MegE progenitor
MPP + CMP
Neutrophil
C
HSC/MPP CLUSTER
WT
KO
HSC
MPP
UMAP 1
UMAP 2
D
HSC SCORE
ACTIVE_HSCs_MPPS
UMAP 1
UMAP 2
E
H19
Meg3
Mki67
CDK1
High
Low
UMAP 1
UMAP 2
F
DORMANCY MODULE
ACTIVITY MODULE
Hlf
Mecom
Procr
Mllt3
Meg3
H19
Cdk4
Cdk6
Myc
Mki67
Cd48
Cdk1
Top2a
Cdc20
Cd34
ACTIVE
DORMANT
INTERMEDIATE
Signatures expression
Sample
WT
KO
Phase
G1
G2M
S
G
Dormant
Serial_FL
Diapause
Lowoutput
Activated_HSC_MPPs
Highoutput
HALLMARK_MYC_V1
HALLMARK_MYC_V2
ACTIVE
INTERMEDIATE
DORMANT
Signatures expression
Sample
WT
KO
Phase
G1
G2M
S
H
WT
NICD
LSK 2,5
LSK 15
CKIT
SCA1
MPP3–4 52
MPP2 15
HSPCs 13
20
MPP3–4 90
MPP2 1,1
HSPCs 0,5
8,8
CD48
CD150
G1 29,8
S 10,8
G2M 21,7
G0 33,8
G1 50,0
S 9,68
G2M 16,1
G0 19,4
Ki67
DAPI
I
NICD vs WT
RNAseq data analysis (23532) genes
lFC cutoff: 0.5; adj p-val cutoff: 0.1
HSC activation markers
HSC markers
Notch target genes
Other
DOWN–REG N=346
UP–REG N=876
Hes1
Hey1
Nrarp
Cd74
Cd69
Itga9
Nfkbia
Mllt3
Cd63
Cdk6
Meg3
Gfi1
Hlf
Hoxa9
–Log10(adj pval)
Shrunken log2 FC

### FIGURE S3

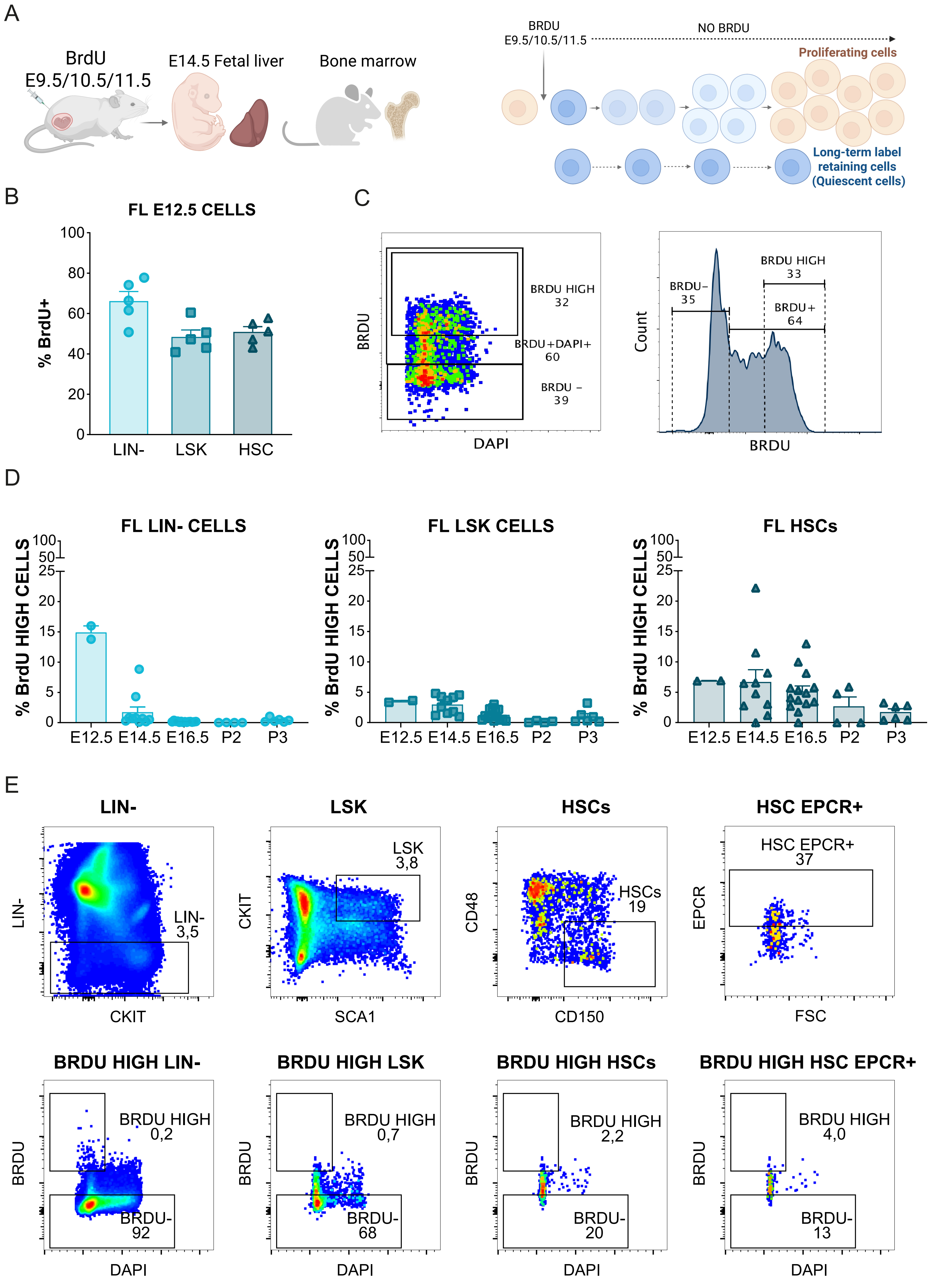
A
BrdU
E9.5/10.5/11.5
E14.5 Fetal liver
Bone marrow
BRDU
E9.5/10.5/11.5
NO BRDU
Proliferating cells
Long-term label retaining cells (Quiescent cells)
B
FL E12.5 CELLS
% BrdU+
LIN-
LSK
HSC
C
BRDU
DAPI
BRDU HIGH 32
BRDU+DAPI+ 60
BRDU − 39
Count
BRDU− 35
BRDU HIGH 33
BRDU+ 64
D
FL LIN- CELLS
FL LSK CELLS
FL HSCs
% BrdU HIGH CELLS
E12.5
E14.5
E16.5
P2
P3
E
LIN-
LSK
HSCs
HSC EPCR+
CKIT
SCA1
CD48
CD150
EPCR
FSC
LIN- 3,5
LSK 3,8
HSCs 19
HSC EPCR+ 37
BRDU HIGH LIN-
BRDU HIGH LSK
BRDU HIGH HSCs
BRDU HIGH HSC EPCR+
BRDU HIGH 0,2
BRDU- 92
BRDU HIGH 0,7
BRDU- 68
BRDU HIGH 2,2
BRDU- 20
BRDU HIGH 4,0
BRDU- 13

### FIGURE S4

A

B

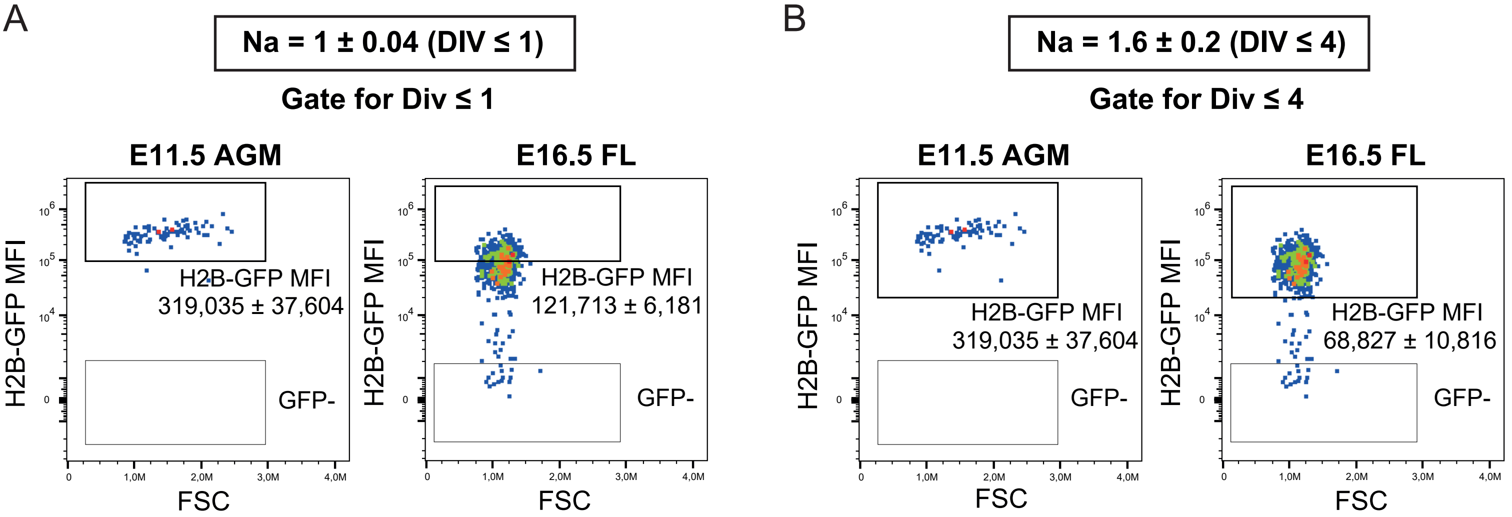


C

| From | To | Na DIVISIONS HSC GFP+ | Na DIVISIONS HSC GFP div≤4 |
|---|---|---|---|
| E11.5 AGM | E16.5 | 1,6 | 1,6 |
| | FL P2 | 3,2 | 2,6 |
| | FL P5 | 5,0 | 3,1 |
| | FL P10 | 3,6 | 2,8 |
| | BM P2 | 4,1 | 3,0 |
| | BM P5 | 14,7* | 2,3 |
| | BM P10 | 9,5* | 3,0 |
| | BM P20 | 12,3* | 3,2 |
| | BM P62 | 19,9* | 3,8 |

*Na accurately estimates up to 6.5 cumulative cell divisions

D

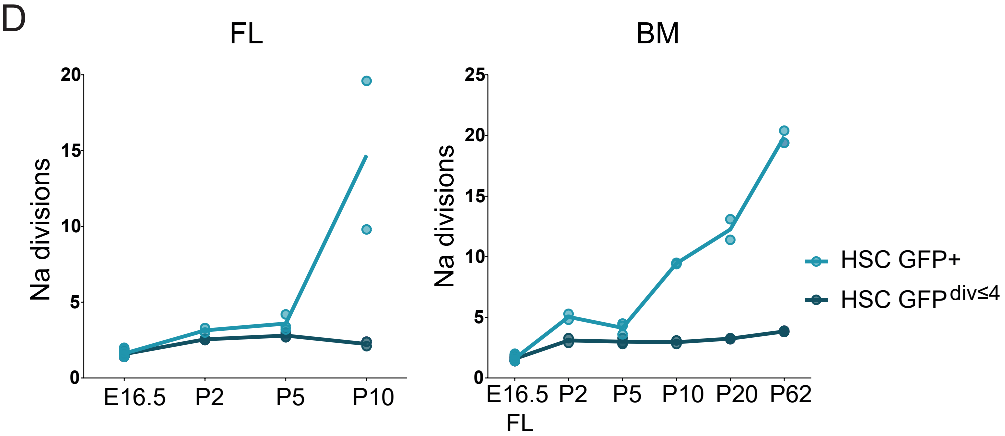


E

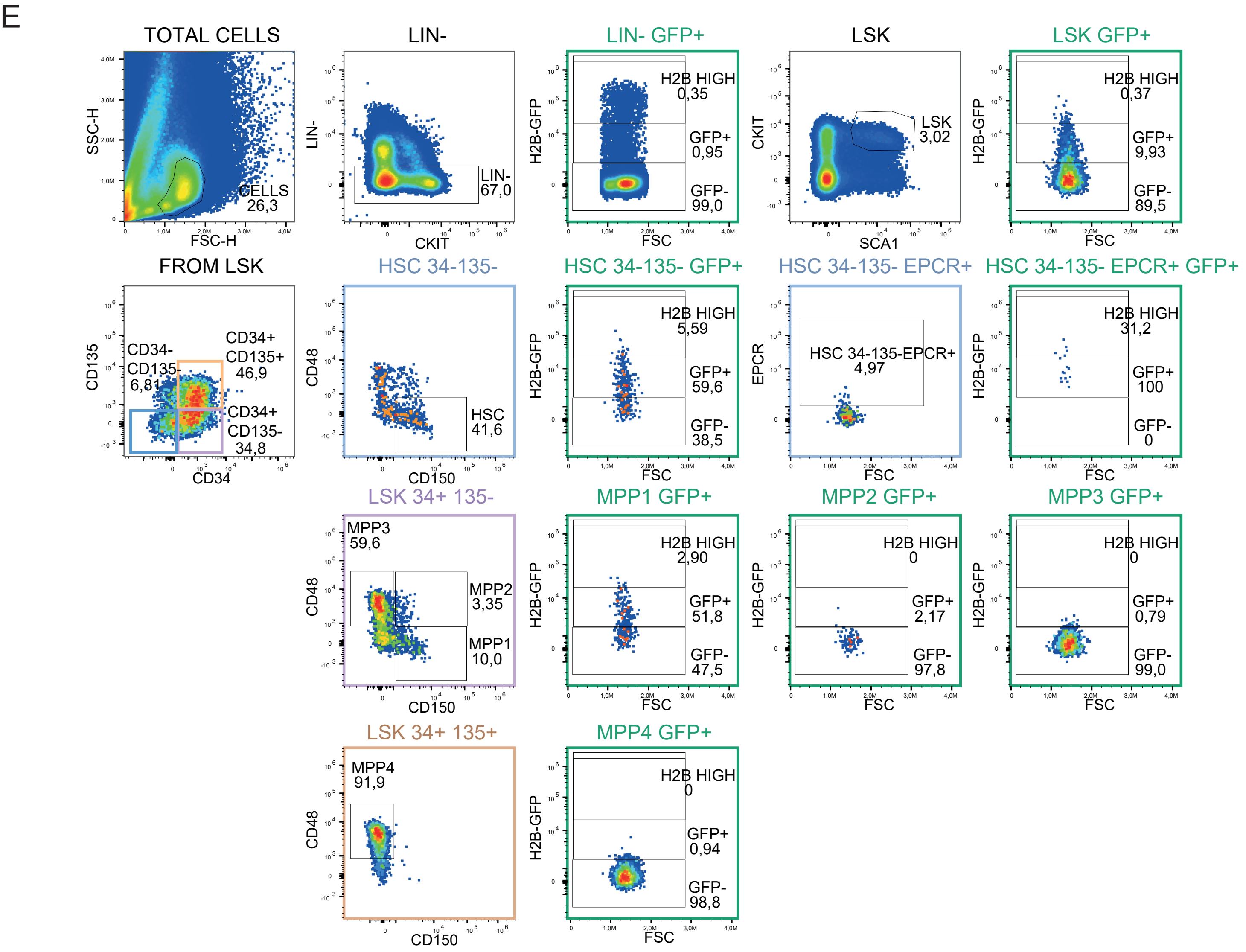

### FIGURE S5

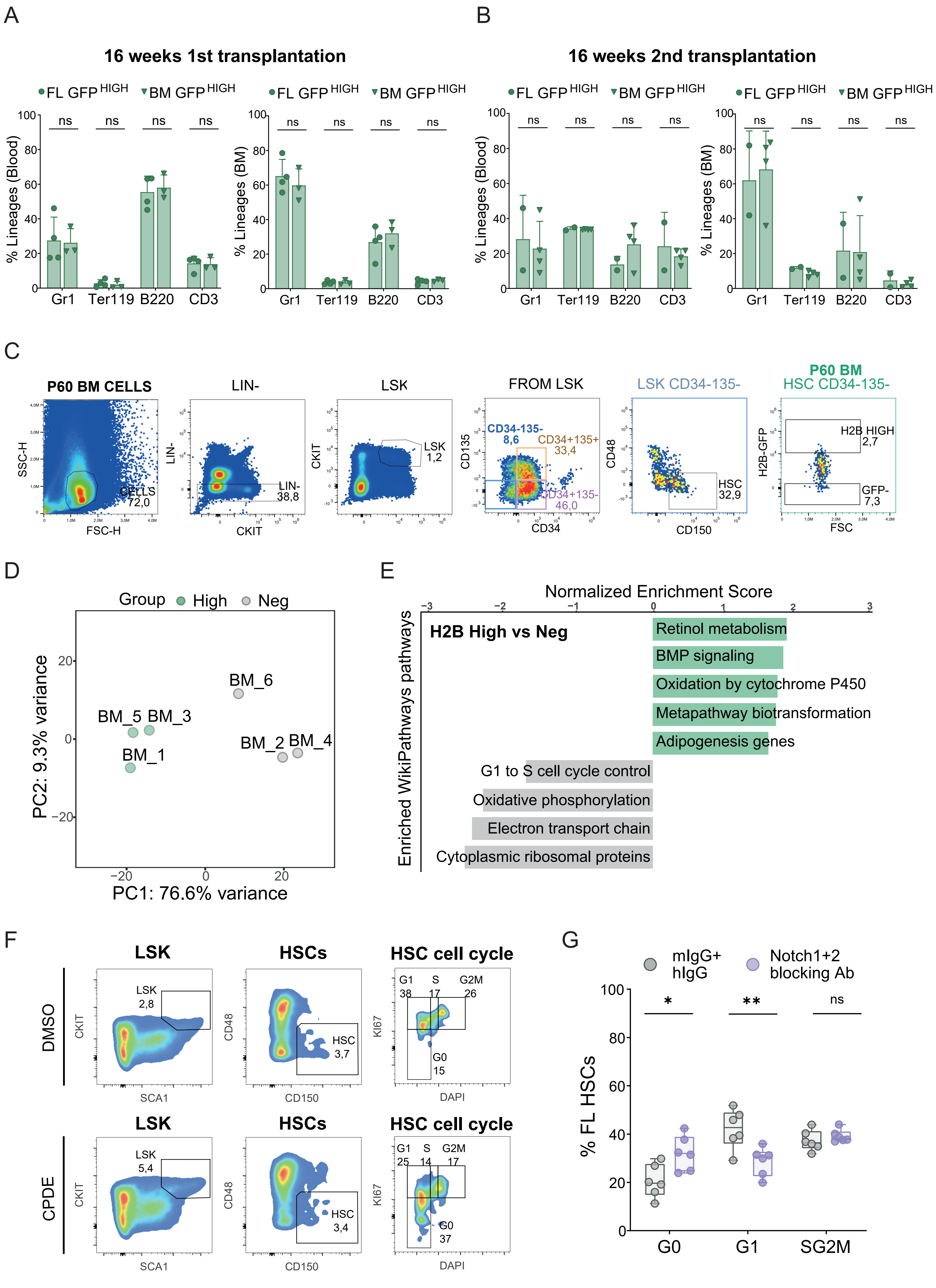
A
B
16 weeks 1st transplantation
16 weeks 2nd transplantation
FL GFP HIGH
BM GFP HIGH
% Lineages (Blood)
% Lineages (BM)
ns
Gr1
Ter119
B220
CD3
C
P60 BM CELLS
LIN-
LSK
FROM LSK
LSK CD34-135-
P60 BM
HSC CD34-135-
SSC-H
FSC-H
CELLS
72,0
LIN-
CKIT
LIN-
38,8
LSK
1,2
CD135
CD34
CD34-135-
8,6
CD34+135+
33,4
CD34+135-
46,0
CD48
CD150
HSC
32,9
H2B-GFP
FSC
H2B HIGH
2,7
GFP-
7,3
D
Group
High
Neg
BM_6
BM_5 BM_3
BM_1
BM_2 BM_4
PC2: 9.3% variance
PC1: 76.6% variance
E
Normalized Enrichment Score
Enriched WikiPathways pathways
H2B High vs Neg
Retinol metabolism
BMP signaling
Oxidation by cytochrome P450
Metapathway biotransformation
Adipogenesis genes
G1 to S cell cycle control
Oxidative phosphorylation
Electron transport chain
Cytoplasmic ribosomal proteins
F
LSK
HSCs
HSC cell cycle
DMSO
CPDE
CKIT
SCA1
LSK
2,8
CD48
CD150
HSC
3,7
KI67
DAPI
G1
38
S
17
G2M
26
G0
15
LSK
5,4
HSC
3,4
G1
25
S
14
G2M
17
G0
37
G
mIgG+
hIgG
Notch1+2
blocking Ab
% FL HSCs
*
**
ns
G0
G1
SG2M

### FIGURE S6

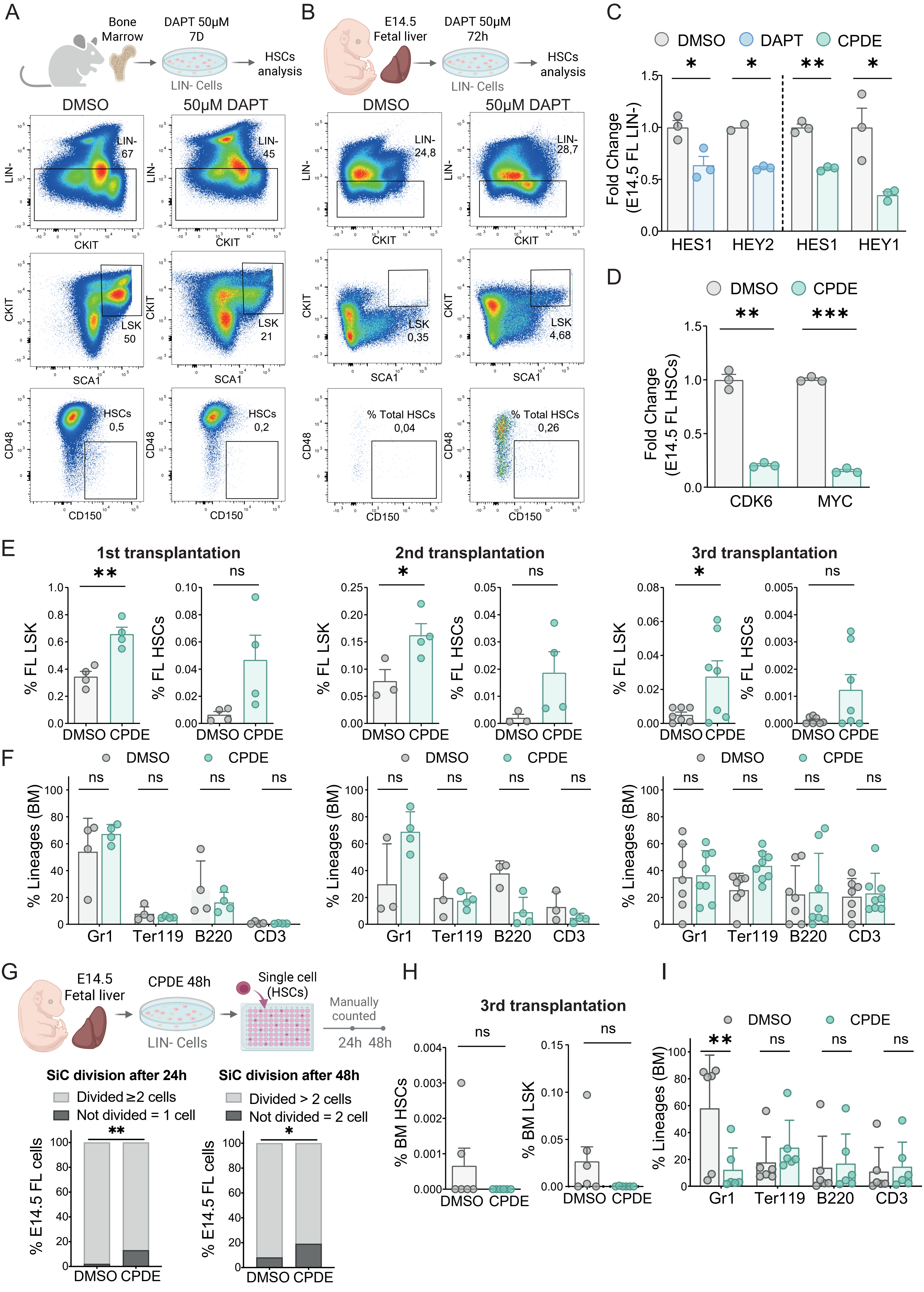

A
Bone Marrow
DAPT 50µM 7D
LIN- Cells
HSCs analysis
DMSO
50µM DAPT
LIN- 67
LIN- 45
LIN-
CKIT
LSK 50
LSK 21
SCA1
HSCs 0,5
HSCs 0,2
CD48
CD150
B
E14.5 Fetal liver
DAPT 50µM 72h
LIN- Cells
HSCs analysis
DMSO
50µM DAPT
LIN- 24,8
LIN- 28,7
LSK 0,35
LSK 4,68
% Total HSCs 0,04
% Total HSCs 0,26
C
DMSO
DAPT
CPDE
Fold Change (E14.5 FL LIN-)
HES1
HEY2
HES1
HEY1
D
DMSO
CPDE
Fold Change (E14.5 FL HSCs)
CDK6
MYC
E
1st transplantation
2nd transplantation
3rd transplantation
% FL LSK
% FL HSCs
DMSO CPDE
F
DMSO
CPDE
% Lineages (BM)
Gr1
Ter119
B220
CD3
G
E14.5 Fetal liver
CPDE 48h
LIN- Cells
Single cell (HSCs)
Manually counted
24h 48h
SiC division after 24h
Divided ≥2 cells
Not divided = 1 cell
SiC division after 48h
Divided > 2 cells
Not divided = 2 cell
% E14.5 FL cells
DMSO CPDE
H
3rd transplantation
% BM HSCs
% BM LSK
DMSO CPDE
I
DMSO
CPDE
% Lineages (BM)
Gr1
Ter119
B220
CD3
